# Structural basis for activation of DNMT1

**DOI:** 10.1101/2022.06.16.496385

**Authors:** Amika Kikuchi, Hiroki Onoda, Kosuke Yamaguchi, Satomi Kori, Shun Matsuzawa, Yoshie Chiba, Shota Tanimoto, Sae Yoshimi, Hiroki Sato, Atsushi Yamagata, Mikako Shirouzu, Naruhiko Adachi, Jafar Sharif, Haruhiko Koseki, Atsuya Nishiyama, Makoto Nakanishi, Pierre-Antoine Defossez, Kyohei Arita

**Affiliations:** Structural Biology Laboratory, Graduate School of Medical Life Science, Yokohama City University, Tsurumi-ku, Yokohama, Kanagawa, 230-0045, Japan; Université Paris Cité, CNRS, Epigenetics and Cell Fate, 75013 Paris, France; Division of Cancer Cell Biology, The Institute of Medical Science, The University of Tokyo, 4-6-1 Shirokanedai, Minato-ku, Tokyo 108-8639, Japan; Laboratory for Protein Functional and Structural Biology, RIKEN Center for Biosystems Dynamics Research, 1-7-22 Suehiro-cho, Tsurumi-ku, Yokohama, Kanagawa 230-0045, Japan; Structural Biology Research Center, Photon Factory, Institute of Materials Structure Science, High Energy Accelerator Research Organization (KEK), Tsukuba, Ibaraki 305-0801, Japan; Laboratory for Developmental Genetics, RIKEN Center for Integrative Medical Sciences (IMS), Yokohama, Kanagawa, Japan

## Abstract

DNMT1 is an essential enzyme that maintains genomic DNA methylation, and its function is regulated by mechanisms that are not yet fully understood. Here, we report the cryo-EM structure of human DNMT1 bound to its two natural activators: hemimethylated DNA and ubiquitinated histone H3. We find that a hitherto unstudied linker, between the RFTS and CXXC domains, plays a key role for activation. It contains a conserved α-helix which engages a crucial "Toggle" pocket, displacing a previously described inhibitory linker, and allowing the DNA recognition helix to spring into the active conformation. This is accompanied by large-scale reorganization of the inhibitory RFTS and CXXC domains, allowing the enzyme to gain full activity. Our results therefore provide a mechanistic basis for the activation of DNMT1, with consequences for basic research and drug design.

## Introduction

DNA methylation is a key epigenetic mark that regulates gene expression and genome stability ^1, 2^. In mammals, DNA methylation occurs at the 5^th^ position of cytosine, mostly within CpG dinucleotides, and it is catalyzed by a DNA methyltransferase (DNMT) family. *De novo* DNMTs (DNMT3A, 3B and 3L) set up proper DNA methylation pattern during development and differentiation, and this pattern is faithfully copied on the newly replicated DNA at each round of cell division, by the maintenance enzyme DNMT1 ^3, 4^. An E3 ubiquitin ligase (ubiquitin-like containing PHD and RING finger domains 1, UHRF1) protein, plays a crucial role for maintenance DNA methylation ^5, 6^, together with DNMT1. UHRF1 recognizes hemimethylated DNA via its SET and RING-associated (SRA) ^7–9^ and catalyzes double monoubiquitination at K18 and K23 on histone H3 (H3Ub2) which, in turn, recruits DNMT1 and stimulates its enzymatic activity ^10–14^.

DNMT1 is a large protein (1616 amino acids), containing multiple domains (Fig. 1a), and subject to intramolecular regulations that strongly restrict its activity to hemimethylated DNA ^15^. In the absence of DNA (apo-DNMT1, PDB:4WXX, aa:351-1600), the enzyme is autoinhibited: Binding of Replication-Foci Targeting Sequence (RFTS) to the catalytic core, in association with recognition of the DNA binding region by an Auto-Inhibitory Linker, inhibit the access of hemimethylated DNA to DNMT1 catalytic region ^16, 17^. A key unresolved question is: how does the combined presence of H3Ub2 and hemimethylated DNA allow the enzyme to overcome this double inhibition? Of note, previous structural studies of DNMT1 in a complex with hemimethylated DNA (PDB:4DA4, aa:731-1602; 6X9I, aa:729-1600) ^18–20^, in a complex with unmethylated CpG DNA (PDB:3PTA, aa:646-1600) ^21^ and RFTS bound to H3Ub2 (PDB:5WVO, aa:351-600; 6PZV, aa:349-594) ^12, 13^ have used a truncated version of the protein (Fig. 1a), therefore the fate of the inhibitory regions, RFTS and Auto-Inhibitory Linker, during activation is unknown.

**Fig. 1.**
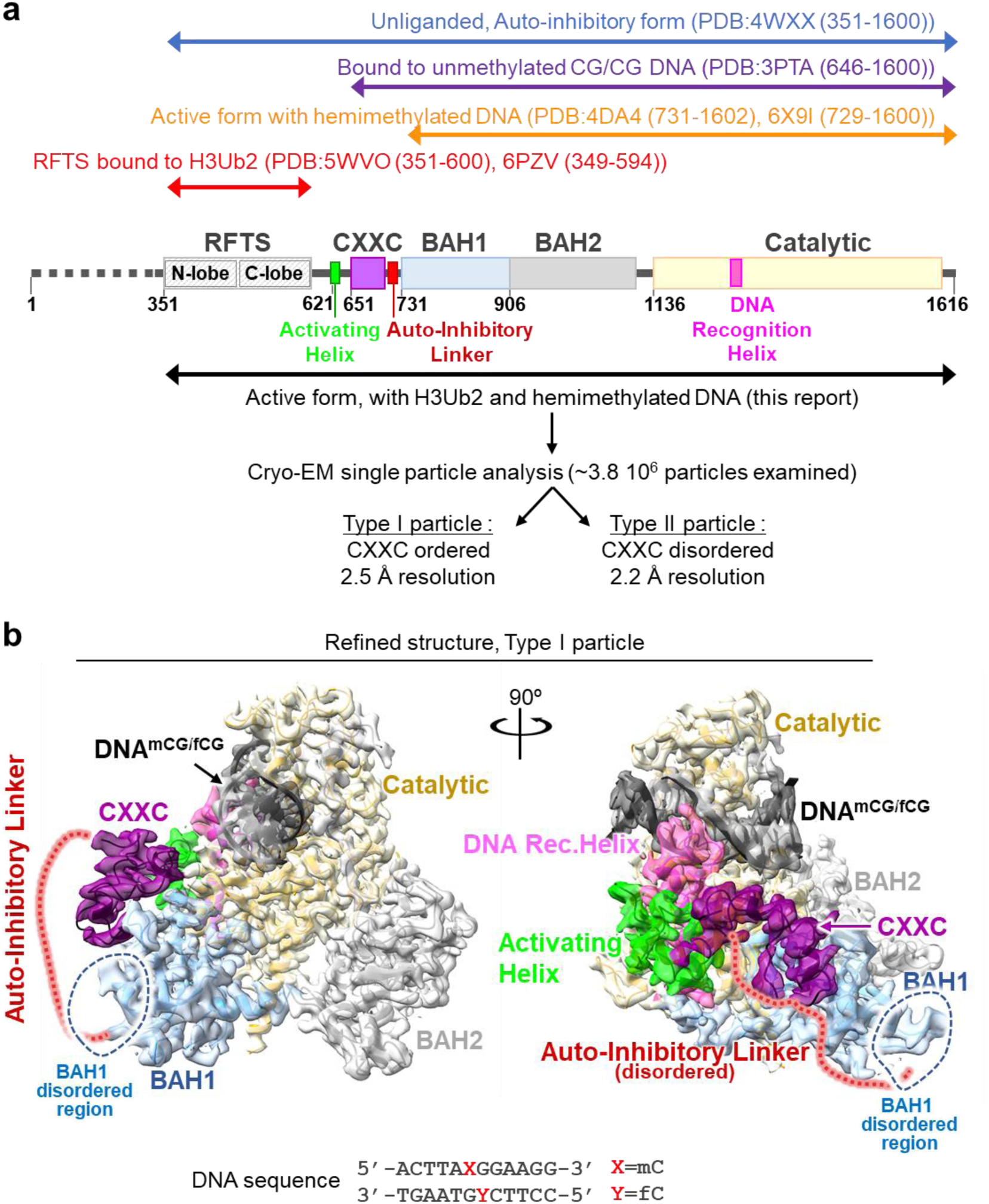
Cryo-EM single particle analysis of DNMT1:H3Ub2:DNA^mCG/fCG^. **a,** Domain architecture of hDNMT1, with amino acid numbers indicated. **b,** Cryo-EM map of the DNMT1:H3Ub2:DNA^mCG/fCG^ ternary complex superposed on the cartoon model. Disordered Auto-Inhibitory Linker and N-terminal portion of BAH1 domain are shown as dotted lines.

In order to understand the detailed molecular mechanism for DNMT1 activation, we have determined the cryogenic electron microscopy (cryo-EM) structure of human DNMT1 (aa:351-1616), stimulated by the H3Ub2 tail and in an intermediate complex with a hemimethylated DNA analog. Our structure illuminates the synergistic structural rearrangements that underpin the activation of DNMT1. In particular, it highlights the key role of a hydrophobic "Toggle" pocket in the catalytic domain, which stabilizes both the inactive (inhibited) or the active states. In the latter case, it functions by accepting a pair of phenylalanine residues from a hitherto unrecognized, yet highly conserved Activating Helix located between the RFTS and CXXC domains.

## Results

### Cryo-EM structure of DNMT1 bound to ubiquitinated H3 and hemimethylated DNA

To uncover the molecular mechanism of DNMT1 activation, we conducted cryo-EM single particle analysis of DNMT1 (aa:351-1616) in an intermediate complex with H3Ub2 tail and hemimethylated DNA (Fig. 1). The human DNMT1 protein, a minimum fragment required for investigating the activation mechanism by binding of ubiquitinated H3, was produced using the Sf9 baculovirus expression system, and an H3 tail peptide (aa:1-37W, K14R/K27R/K36R) was di-ubiquitinated on K18 and K23 to completion *in vitro* (Extended Data Fig. 1 and see method). As expected from previous work ^12, 13^, in contrast to K18 or K23 single mono-ubiquitinated H3, the addition of H3Ub2 effectively enhanced the enzymatic activity of DNMT1 (Extended Data Fig. 1d,e). The DNMT1:H3Ub2 binary complex was used for reaction with hemimethylated DNA. The target cytosine in hemimethylated DNA was replaced by a 5-fluorocytosine (5fC) to form an irreversible covalent complex with DNMT1 ^22^. The ternary complex containing DNMT1 bound to H3Ub2 and DNA^mCG/fCG^ was purified by gel-filtration chromatography (Extended Data Fig. 2), and used for cryo-EM single particle analysis (Extended Data Fig. 3).

3D variability analysis by cryoSPARC ^23^ revealed 2 types of particles: Type I particles with 2.5 Å resolution, in which the CXXC domain was ordered (Fig. 1b and Extended Data Fig. 4a), and Type II particles with 2.2 Å resolution, in which the CXXC was disordered (Extended Data Fig. 5a). The atomic models of DNMT1 were constructed from the two types (Table 1); these structures were essentially the same (except for the CXXC domain), therefore in the rest of the paper we will focus on the Type I DNMT1 particle, with the ordered CXXC. In this structure, the RFTS domain, Auto-Inhibitory Linker, N-terminal β-sheet of BAH1 (aa:731-755) and some loops and linkers were invisible, reflecting their flexibility (Fig. 1b). The structured elements of the ternary complex showed that the catalytic core of DNMT1 bound to hemimethylated DNA, the catalytic loop (aa:1224-1238) recognized flipped-out 5fC, and the Target Recognition Domain (TRD) residues (Cys1499, Leu1500, Trp1510, Leu1513 and Met1533) bound to the methyl-group of 5mC (Extended Data Fig. 5b-d). Overall, these results confirm previously reported DNMT1:hemimethylated DNA binary complex structures (Supplementary Notes) ^18–20^. In addition, they show, for the first time, the behavior of the RFTS and Auto-Inhibitory Linker in the active form, as described in the following section.

### Spatial rearrangement of RFTS and CXXC domains in the active form

The dissociation of the RFTS domain from the catalytic core is assumed to be required for DNMT1 activation, yet no direct evidence of this structural rearrangement has been shown to date. Our cryo-EM structure of apo-DNMT1 (aa:351-1616) at 3.4 Å resolution showed a fully-structured RFTS domain bound to the catalytic core (Extended Data Fig. 6a left, and Supplementary Table 1). In contrast, the cryo-EM map of the DNMT1:H3Ub2:DNA^mCG/fCG^ ternary complex showed no density for the RFTS:H3Ub2 (Fig. 1b). Intriguingly, particles smaller than the DNMT1:H3Ub2:DNA^mCG/fCG^ ternary complex were observed in 2D class average (Extended Data Fig. 7a). The size and shape of these particles were comparable to those of RFTS:H3Ub2 as estimated from the 2D projected template of RFTS:H3Ub2 (PDB:5WVO)-derived 3D Gaussian model (Extended Data Fig. 7b). These data indicate that the RFTS:H3Ub2 moiety is in a highly dynamic state and does not interact with other domains when DNMT1 is active.

To separate the contributions of H3Ub2 and of hemimethylated DNA for displacement of the RFTS, we determined the cryo-EM structure of DNMT1 (aa:351-1616) in complex with H3Ub2, but without DNA (Extended Data Figs. 1c, 6a right, and Supplementary Table 1). This structure reached 3.6 Å resolution and showed that, while the N-lobe of DNMT1 became flexible and therefore invisible, the C-lobe remained bound firmly to the catalytic core upon H3Ub2 binding. These data were further supported by small angle X-ray scattering (SAXS) analyses (Extended Data Fig. 6b, Supplementary Fig. 6, Supplementary Table 2, and Supplementary Notes), showing that H3Ub2, in itself, is not sufficient to dislodge the RFTS.

We next investigated the dynamics of the CXXC domain after DNMT1 activation. In apo-DNMT1, the CXXC domain is affixed to the side of the RFTS domain (Fig. 2a) ^16, 17^. In the presence of unmethylated DNA (DNA^CG/CG^) (PDB:3PTA), the zinc finger motif of the CXXC domain recognizes unmethylated CpG, shifts 30 Å towards the TRD domain, and sits near the active center (Fig. 2a) ^21^. In this conformation, the Auto-Inhibitory Linker directly interrupts the binding of DNA to the active site, which prevents unlicensed *de novo* DNA methylation. Intriguingly, in our active complex, the CXXC domain moved away from the active center and took an "upside-down" position (relative to 3PTA), between the BAH1 domain and the catalytic domain with which it established hydrogen bonds and van der Waals interactions (Fig. 2b,c).

**Fig. 2.**
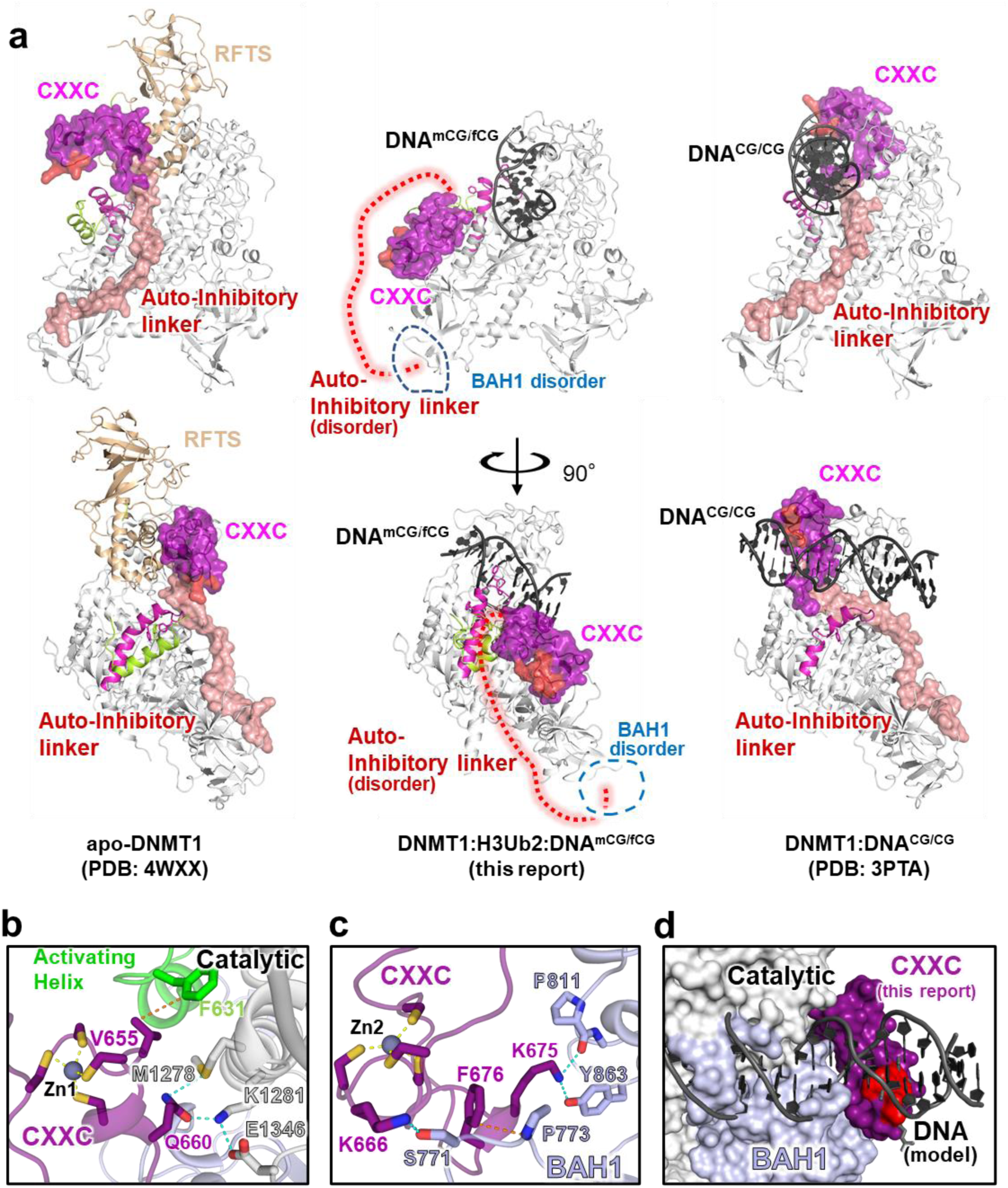
Spatial rearrangement of the CXXC domain upon activation. **a,** Comparison of overall structure of apo-DNMT1 (left, PDB:4WXX), DNMT1:H3Ub2:DNA^mCG/fCG^ ternary complex (center, this study), and DNMT1:DNA^CG/CG^ binary complex (right, PDB: 3PTA). Upper and lower panels show the side and front views of DNMT1, respectively. RFTS domain, Activating Helix, CXXC domain, Auto-Inhibitory Linker, and DNA Recognition Helix were colored light orange, light green, purple, red and pink, respectively. CXXC domain and Auto-Inhibitory Linker are also exhibited as the transparent surface model. The disordered regions of BAH1 and Auto-Inhibitory Linker in DNMT1:H3Ub2:DNA^mCG/fCG^ ternary complex are shown as dotted line. The red surface in the CXXC domain indicates R681-K683, which bind the major groove of unmethylated DNA^CG/CG^. **b,** Detail of the interaction between CXXC domain and catalytic domain showing gray cartoon and stick models in the ternary complex. **c,** Detail of the interaction between CXXC domain and BAH1 domain showing light-purple cartoon and stick models in the ternary complex. **d,** Model structure of CXXC domain bound to DNA^CG/CG^ in the ternary complex. CXXC:DNA^CG/CG^ in PDB:3PTA is superimposed on the CXXC domain in the ternary complex. The red surface in the CXXC domain indicates the binding surface for binding to the major groove of unmethylated DNA^CG/CG^.

As the DNA binding interface of the CXXC remained solvent-exposed in the active complex (Fig. 2a), so we asked whether it could still bind unmethylated DNA. However, the superimposition of the CXXC:DNA complex onto the ternary complex structure revealed major steric clashes with the catalytic domain (Fig. 2d), suggesting that DNA binding by the CXXC is fully suppressed when DNMT1 is active. The new position of the CXXC leads to a structural deformation of the BAH1 N-terminal β-sheet and induces a different orientation of the BAH1 loop (aa:765-775) (Fig. 1b). These structural changes then lead to full eviction of the Auto-Inhibitory Linker from the catalytic core and causes the Auto-Inhibitory Linker to adopt a highly disordered structure (Figs. 1b and 2a).

Taken together, our cryo-EM analysis of the active form of DNMT1 revealed a large reorganization of the inhibitory domains relative to unliganded or inactive forms. Upon joint binding of H3Ub2 and hemimethylated DNA, the RFTS domain is forced out of the catalytic core, while the CXXC domain undergoes a drastic spatial rearrangement ultimately leading to eviction of the Auto-Inhibitory Linker from the catalytic core.

### A "Toggle" pocket accepts different phenylalanines in the repressed and active states

Zooming in on the catalytic site, we observed another striking difference between the inactive and active states of the enzyme.

In the inactive state, the DNA Recognition Helix (aa:1236-1259) in the catalytic domain is kinked, at Ser1246 (Fig. 3b). A hydrophobic pocket (composed of Phe1229, Val1248, Phe1263, Leu1265, Phe1274, Val1279, and Leu1282, hereafter Toggle Pocket) accepts Phe1235 of the catalytic loop and Phe1243 of the DNA Recognition Helix (Fig. 3a). In addition, Tyr1240 within this same helix forms hydrophobic interactions with Phe628, Phe631 and Phe632 from the Activating Helix (aa:620-635), further stabilizing the inactive conformation (Fig. 3b).

**Fig. 3.**
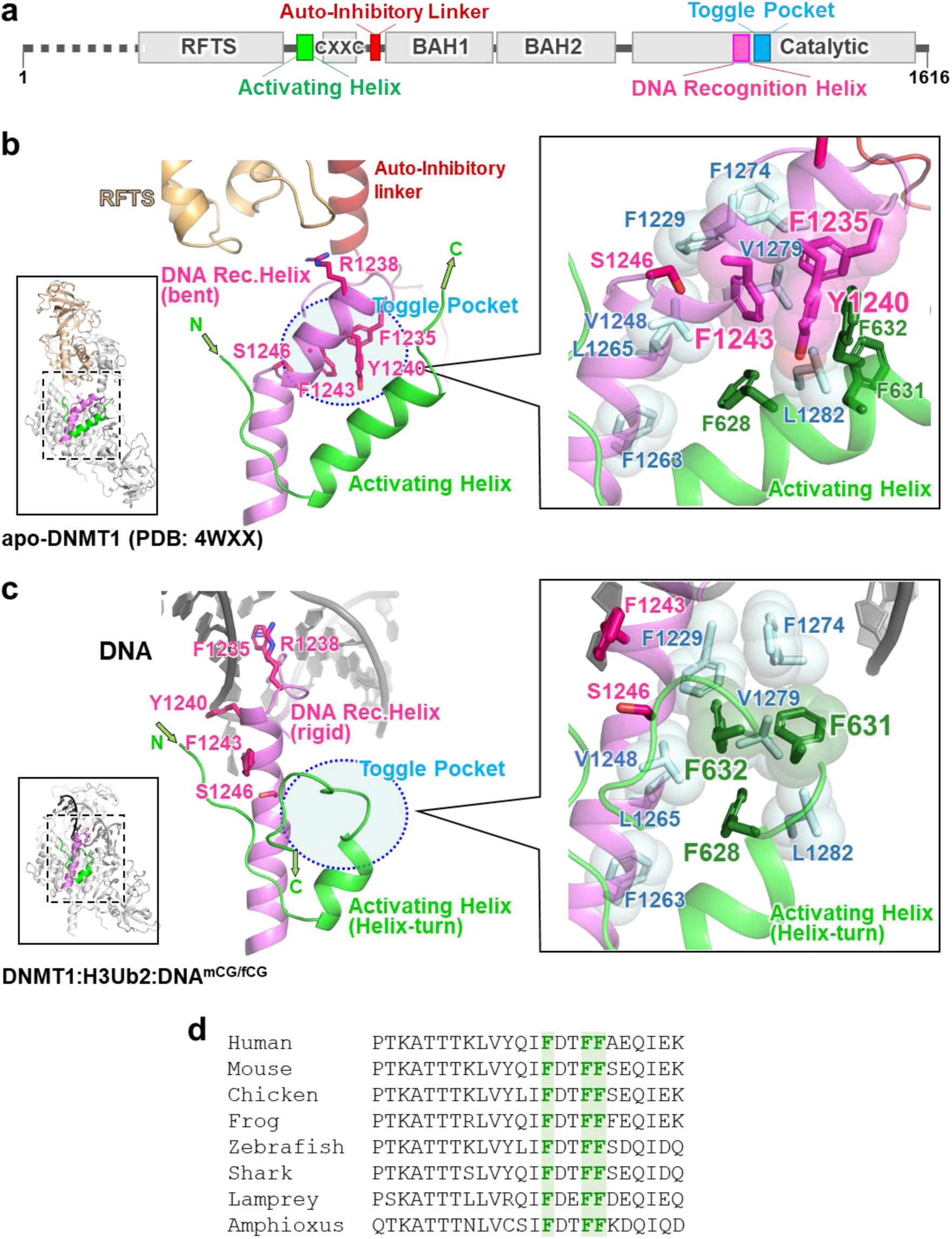
Activation of DNMT1 by a pair of phenylalanine residues in the Activating Helix. **a,** Domain structure of DNMT1. **b,** Structure around Activating Helix (green), DNA Recognition Helix (pink) and Toggle Pocket (pale cyan) of apo-DNMT1 (PDB:4WXX). The Toggle Pocket is highlighted by a light blue circle. Right panel is magnified figure of the Toggle Pocket. Residues in Activating Helix and DNA Recognition Helix that are involved in binding to the Toggle Pocket are shown as green and pink stick models, respectively. Residues for formation of the Toggle Pocket are shown as stick with transparent sphere model. **c,** Structure around Activating Helix (green), DNA Recognition Helix (pink) and Toggle Pocket (pale cyan) of the ternary complex. Color scheme is same as Fig. 3b. **d,** Multiple sequence alignment of DNMT1 around the Activating Helix. Phenylalanines 628, 631 and 632 are highlighted.

This contrasts sharply with our activated form of DNMT1 (Fig. 3c). In that situation, the Activating Helix is shortened as it forms a helix-turn; its residues Phe631 and Phe632 invade the Toggle Pocket. The DNA Recognition Helix is freed from the Toggle Pocket and springs into a straight conformation, which allows i) access of Phe1235/Arg1238 (in the catalytic loop) to the minor groove at the mCG/fCG site, ii) formation of a hydrogen bond between Tyr1240 in the DNA Recognition Helix and the phosphate backbone of the DNA and iii) engagement of Phe1243 (in the DNA Recognition Helix) by Pro613, Lys617 and Gln635 (in the linker between RFTS and CXXC domains), which prevents Phe1243 from entering the Toggle Pocket (Fig. 3a,b and Extended Data Fig. 8a).

Interestingly, a previous study of human DNMT1 binary complex (aa:729-1600, not containing Activating Helix) with hemimethylated DNA analog (active state, PDB: 6X9I) shows that Phe1243 remains in the Toggle Pocket and the N-terminal region of the DNA Recognition Helix structure is partially unfolded, thereby preventing the binding of Tyr1240 to the phosphate backbone of DNA (Extended Data Fig. 8a,b) ^20^. Furthermore, folded or unfolded DNA Recognition Helix structures are observed in the previous structures of mouse DNMT1 (aa:731-1602) bound to hemimethylated DNA analog, depending on the sequence around the mCG/fCG ^18, 19^, suggesting that the DNA Recognition Helix is intrinsically flexible; in contrast, our DNMT1:H3Ub2:DNA^mCG/fCG^ complex shows a rigid conformation of the DNA Recognition Helix, indicating that the phenylalanine pair in the Activating Helix contributes to activation state of DNMT1.

### Crucial role of a phenylalanine pair for activation of DNMT1

The phenylalanines Phe631 and Phe632 are invariant between vertebrate species, and are also present in the cephalochordate Amphioxus (Fig. 3d). We therefore asked whether these residues played a role in the activation of DNMT1, as could be expected from the fact that they bind the Toggle Pocket. In a binding assay, we found that the F631A/F632A mutations abolished the ability of DNMT1 to bind hemimethylated DNA (Fig. 4a), even though the H3Ub2-binding ability of DNMT1 was unaffected (Extended Data Fig. 1f). We carried out an *in vitro* DNA methylation assay and, again, observed that the F631A/F632A mutation led to severe defects in DNA methylation (Fig. 4b).

**Fig. 4.**
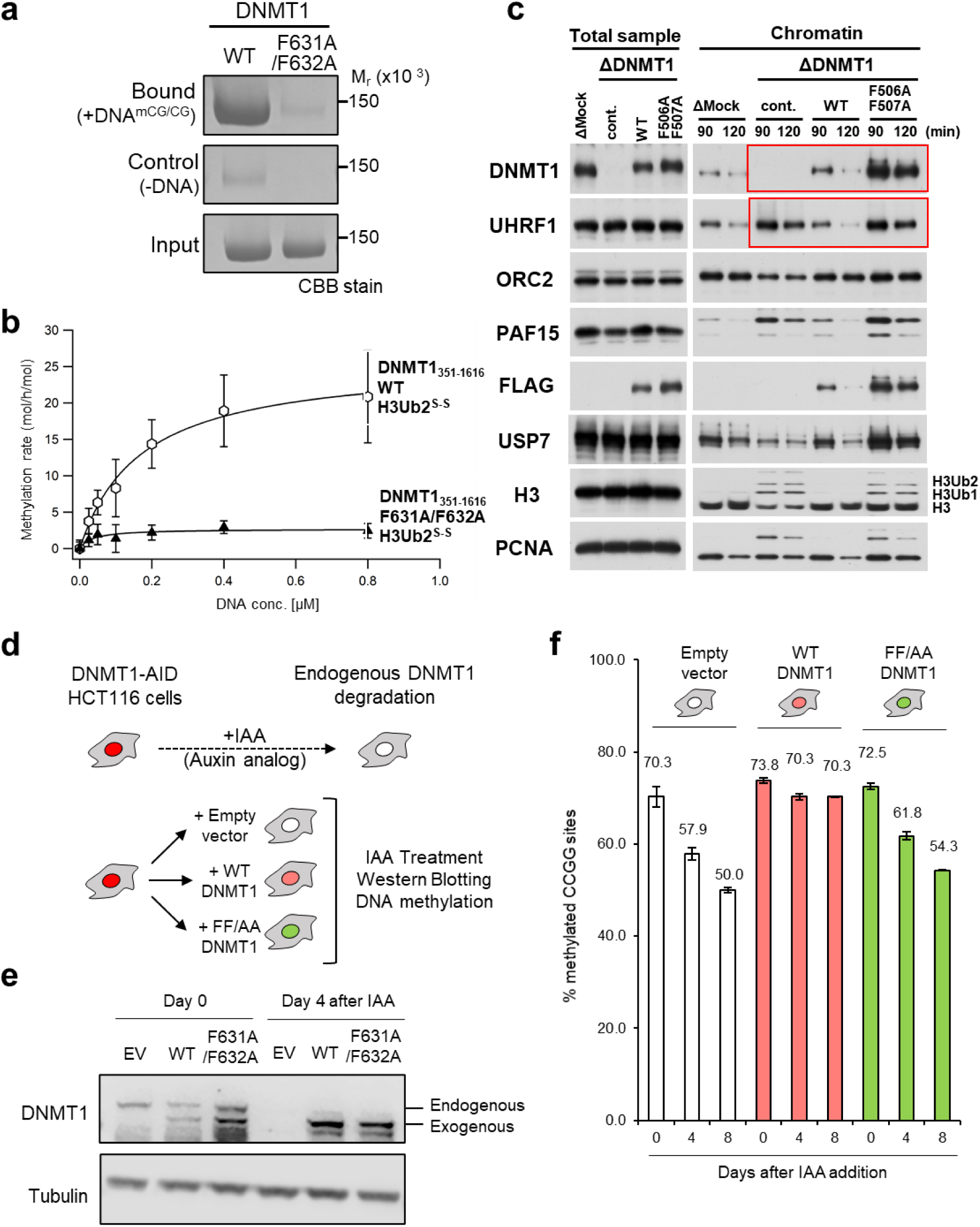
Functional assays demonstrate the importance of the Activating Helix. **a,** Pull-down assay using immobilized hemimethylated DNA. **b,** *in vitro* DNA methylation activity of DNMT1 WT (white hexagons) and F631A/F632A mutant (blacks triangle) for hemimethylated DNA. Vertical axis indicates the turnover frequency of the methylation reaction of DNMT1 (15 nM) after 1 hr. Michaelis–Menten curve is shown as lines. **c,** Interphase extracts depleted with either control or xDnmt1 antibodies were incubated with control buffer, purified recombinant WT xDnmt1 and F506A/F507A. The chromatin fractions were isolated at the indicated times and their bound proteins as well as inputs were analyzed by immunoblotting. **d,** Experimental scheme in human HCT116 colon cancer cells: the endogenous DNMT1 protein is tagged with an Auxin-Inducible Degron (AID) tag, causing the protein to be degraded after addition of the auxin analog IAA. In this background, rescue vectors are added that encode either WT or FF/AA versions of DNMT1. The empty Vector is used as a negative control. **e,** Western blotting shows that endogenous DNMT1 is degraded upon IAA addition, whereas exogenous WT and FF/AA DNMT1 are not degraded. **f,** Measurement of global DNA levels by Luminometric Methylation Assay (LUMA). Cells expressing WT DNMT1 maintain global DNA methylation when endogenous DNMT1 is degraded, whereas the FF/AA mutant does not support DNA methylation maintenance.

We then sought confirmation of these results in an *in vitro* assay, which reconstitutes replication-coupled maintenance DNA methylation using *Xenopus* egg extracts ^11, 12, 24^. In that system, we immunodepleted DNMT1, and re-introduced recombinant DNMT1, either WT or mutated on the 2 phenylalanines of the Activating Helix (F506A and F507A in *Xenopus*, FF/AA mutant). As previously reported, the depletion of xDNMT1 from *Xenopus* egg extracts resulted in the accumulation of chromatin-bound UHRF1 and ubiquitinated histone H3 species (Fig. 4c); this is due to defective maintenance DNA methylation, which generates hemimethylated DNA from which UHRF1 cannot be released ^11, 24^. The addition of wild-type (WT) recombinant xDNMT1 suppressed the accumulation of UHRF1 and ubiquitinated H3 (Fig. 4c). The FF/AA mutant retained chromatin binding activity but failed to suppress the accumulation of UHRF1 and ubiquitinated H3, showing defects in maintenance DNA methylation (Fig. 4c). Therefore, this functional assay in *Xenopus* egg extracts further validated the effect of the mutation.

Lastly, we used a colon cancer cell line HCT116, in which DNA methylation has been widely studied. In this line, both endogenous alleles of DNMT1 are tagged with an Auxin-Inducible Degron (AID) (Fig. 4d). In this DNMT1-AID line, we introduced rescue vectors: one encoding WT DNMT1, and the other encoding the FF/AA mutant (Fig. 4d). The level of endogenous DNMT1, exogenous WT or FF/AA DNMT1 were comparable (Fig. 4e) and they were located in nucleus (Extended Data Fig. 9). Treating the cells with indole-3-acetic acid (IAA) caused the disappearance of endogenous DNMT1 but did not affect the exogenous proteins. We then measured global DNA methylation at days 0, 4, and 8 after endogenous DNMT1 removal (Fig. 4f). The control cells (empty vector) lost almost one-third of total DNA methylation, going from 70% to 50% methylation. This loss was completely prevented by the WT DNMT1 transgene. In contrast, the FF/AA DNMT1 mutant was incapable of sustaining DNA methylation maintenance, and showed DNA methylation values close to those of the empty vector (Fig. 4f).

Collectively these experiments confirm that the Activating Helix, and especially its conserved phenylalanines, are crucial for DNA methylation maintenance by DNMT1.

## Discussion

Our cryo-EM analysis reveals a molecular mechanism for human DNMT1 catalytic activation (Fig. 5). Our results reveal both the large-scale displacements of inhibitory modules (RFTS, CXXC, Auto-Inhibitory Linker), as well as more detailed changes, particularly the switch by which the same hydrophobic pocket, initially bound to inhibitory phenylalanines, engages activating phenylalanines, which releases the DNA Recognition Helix and permits catalysis. This regulation also operates in *Xenopus*, and may even occur in invertebrates such as *Amphioxus*, in which the regulatory amino acids are conserved (Fig. 3d).

**Fig. 5.**
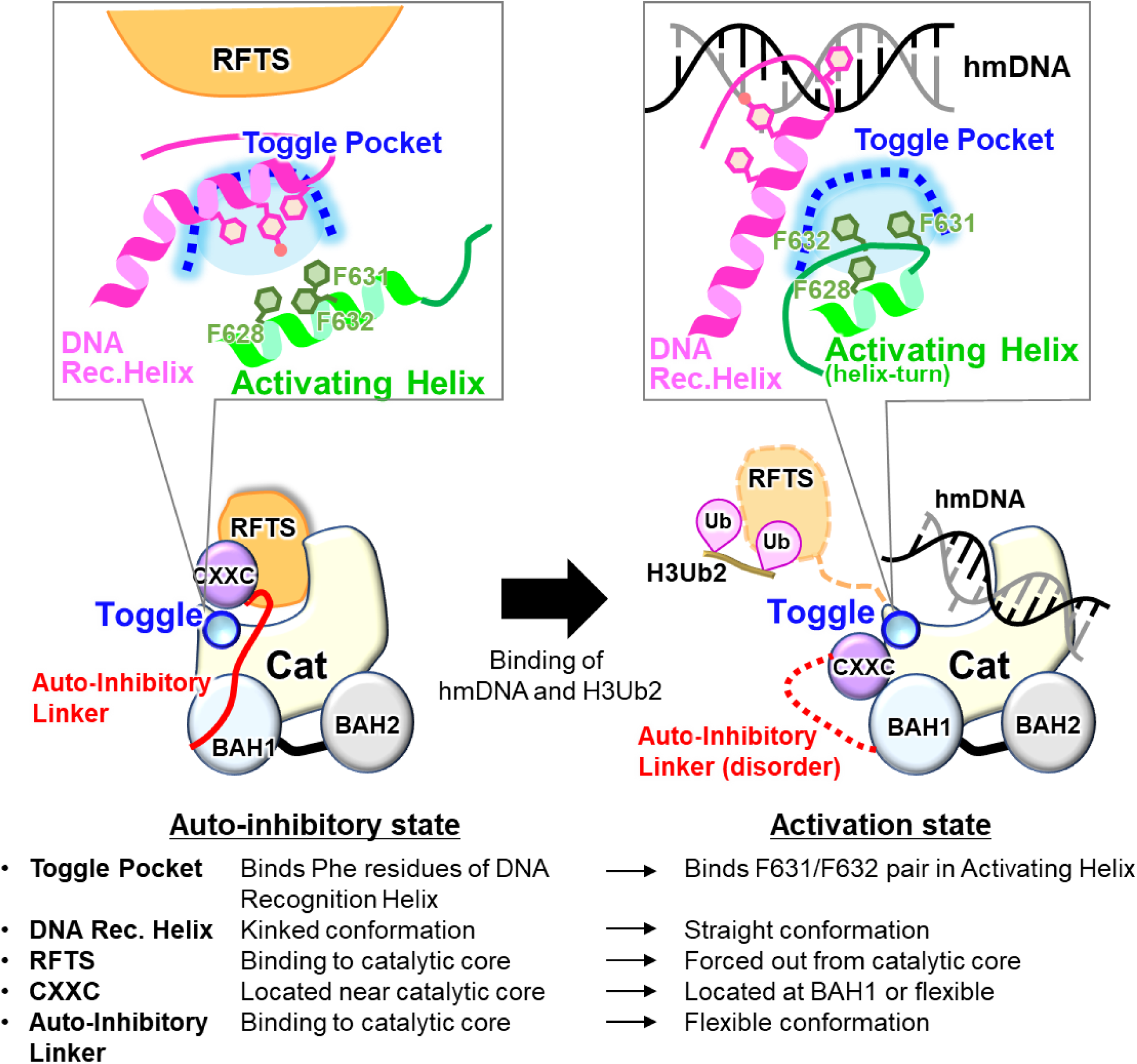
Schematic representation of the DNMT1 activation mechanism. Left and right Figures show the auto-inhibitory and activation state of DNMT1, respectively.

The catalytic domains of the *de novo* DNA methyltransferases DNMT3A and DNMT3B bind the DNMT3L catalytic-like domain and form a heterotetramer ^25–28^. Interestingly, the DNMT3A(B)/3L interface is formed by hydrophobic interactions mediated by phenylalanine residues, and therefore is known as the F-F interface. The F-F interface enhances DNA methylation activity by the DNMT3A(B)/3L heterotetramer ^25^. The hydrophobic residue in the DNMT3A(B) catalytic domain spatially corresponds to the Toggle Pocket of DNMT1. Thus, covering the hydrophobic pocket of the catalytic domain by an intra- or inter-molecule interaction could be an evolutionarily conserved activation mechanism of DNA methyltransferases. The Activating Helix, however, is unique to DNMT1 and crucial for enzymatic activation, and therefore could be utilized to design novel inhibitors such as helical peptides that mimic this Activating Helix.

Previously reported structures of apo-DNMT1 and DNMT1:DNA^CG/CG^ revealed a dual-autoinhibitory mechanism in which the RFTS domain and the Auto-Inhibitory Linker are embedded into the catalytic core, thereby inhibiting the access of cognate DNA (Fig. 5). Our cryo-EM analysis of the ternary complex showed full dissociation of both the RFTS domain and Auto-Inhibitory Linker from the catalytic core (Fig. 1b). Interestingly, H3Ub2-binding to the RFTS domain might not be sufficient for displacement of the RFTS domain as our cryo-EM and SAXS data showed that the C-lobe of RFTS domain is still accommodated in the catalytic core in the DNMT1:H3Ub2 complex (Extended Data Fig. 6). A previous molecular dynamics simulation has demonstrated that H3Ub2-binding reduces the contact number between the C-lobe and catalytic core ^12^. In addition, apo-DNMT1 was unable to form the binary complex with hemimethylated DNA (Extended Data Fig. 1c) We hypothesize, therefore, that H3Ub2 binding destabilizes the inhibitory interaction between the C-lobe and the catalytic core, allowing hemimethylated DNA to penetrate the catalytic core. Thus, we propose that simultaneous binding to H3Ub2 and DNA leads to full activation of DNMT1 via the following structural changes: i) dissociation of the RFTS domain from the catalytic core, ii) structural changes to a newly identified Activating Helix causing the conserved residues Phe631/Phe632 to invade the Toggle Pocket of the catalytic domain, iii) adoption of a rigid conformation by the DNA Recognition Helix, iv) spatial rearrangement of the CXXC domain and v) eviction of the Auto-Inhibitory Linker from the catalytic domain (Fig. 5). However, it is currently unknown how simultaneous binding of H3Ub2 and hemimethylated DNA causes a conformational change in the Activation Helix to place F631/F631 in the Toggle Pocket. Future work, such as molecular dynamics simulation, will determine if these structural changes occur sequentially or simultaneously. Thus, our findings describe new concepts and mechanisms in the multi-step activation process of DNMT1 that ensures faithful maintenance of DNA methylation.

## Methods

### Oligonucleotides

12 base of oligonucleotides (upper: 5’-ACTTA(5mC)GGAAGG, lower: 5’-CCTTC(5fC)GTAAGT) for cryo-EM single particle analysis, 42 base of oligonucleotides (upper: 5’-GGACATC(5mC)GTGAGATCGGAGGC(5mC)GCCTGCTGCAATC(5mC)GGTAG, lower: 5’-CTACCGGATTGCAGCAGGCGGCCTCCGATCTCACGGATGTCC) for DNA methylation assay and 21 base of oligonucleotides (upper: 5’-CAGGCAATC(5mC)GGTAGATCGCA, lower: 5’-biotin-TTGCGATCTACCGGATTGCCTG) for DNA pull-down assay were synthesized by GeneDesign, *Inc*. (Osaka, Japan). 5mC and 5fC mean 5-methylcytosine and 5-fluorocytosine, respectively. To prepare DNA duplex, the mixture of equimolar of complementary oligonucleotides were heated at 95 °C for 2 min and annealed at 4 °C for overnight.

### Protein expression and purification

The gene encoding wild type and mutant of human DNMT1 (residues 351-1616) containing N-terminal ten histidine tag (His-tag) and human rhinovirus 3C (HRV 3C) protease site was amplified by PCR and cloned into the pFastBac vector (Invitrogen) using seamless cloning method. Baculoviruses for expression of the DNMT1 were generated in *Spodoptera frugiperda* 9 (Sf9) cells according to the Bac-to-Bac system instruction (Invitrogen). The protein expression was performed by infection of the baculoviruses with the Sf9 cells for 72 hrs at 27 °C. Cells cultured with 1 L medium were lysed by lysis buffer (50 mM Tris-HCl [pH 8.0] containing 500 mM NaCl, 25 mM Imidazole, 10% Glycerol and 1mM DTT) and sonicated with the cycle of pulse on for 10 sec and pulse off for 50 sec (total pulse on time: 6 min). A soluble fraction was obtained after centrifugation of the lysate at 19,000 rpm for 40 min at 4 °C performed using Avanti J-E with a rotor JA-20 (BECKMAN COULTER) to remove insoluble debris.

The His-tagged DNMT1 was loaded to Ni-Sepharose 6 Fast Flow (Cytiva), and unbound proteins were washed by wash buffer (50 mM Tris-HCl [pH 8.0] containing 1 M NaCl, 25 mM Imidazole, 10% Glycerol and 1 mM DTT), and lysis buffer. Bound proteins were eluted by an elution buffer (20 mM Tris-HCl [pH 8.0] containing 500 mM Imidazole, 300 mM NaCl, 10% Glycerol and 1 mM DTT). The His-tag was cleaved by the HRV3C protease at 4 °C for over 12 hrs. The DNMT1 was separated by anion-exchange chromatography, HiTrap Q HP (Cytiva) using gradient elution from 50 to 1000 mM NaCl in 20 mM Tris-HCl [pH 8.0] buffer containing 10% Glycerol and 1 mM DTT. At the final stage, the DNMT1 was purified with Hiload 26/600 Superdex 200 size exclusion chromatography (Cytiva) equilibrated with 20 mM Tris-HCl [pH 7.5], 250 mM NaCl, 10% Glycerol and 5 mM DTT.

### Preparation of disulfide- and isopeptide-linked ubiquitinated H3

Disulfide-linked ubiquitinated H3 analog for SEC-SAXS and *in vitro* DNA methylation assay was prepared using G76C mutant of ubiquitin (Ub-G76C) and K18C/K23C mutant of H3 peptide (residues 1-36 with an additional tryptophan residue at their C-terminus, hereafter H3_1-37W_-K18C/K23C). Expression in *E. coli* and purification of the Ub-G76C and H3_1-37W_ were purified according to the previous report ^12^. After purification, these proteins were lyophilized. The lyophilized Ub-G76C was dissolved in 50 mM sodium phosphate (pH 7.5) and mixed with a 20-fold molar excess of 5,5′-dithiobis-(2-nitrobenzoic acid) (DTNB, Wako) and the mixture was incubated for 40 minutes at room temperature with rotation on a ROTATOR RT-5 (TITEC). The reaction solution was buffer-exchanged into ligation buffer (20 mM Tris-HCl [pH 7.0], 50 mM NaCl and 1 mM EDTA) using a PD-10 desalting column (Cytiva). Lyophilized H3_1-37W_-K18C/K23C was reduced in 20 mM Tris-HCl (H 7.5) containing 5 mM DTT, and were buffer-exchanged into ligation buffer, and mixed with a 5-fold molar excess of activated Ub-G76C-DTNB for 1 hr. The reaction product was purified on a cation-exchange column, Mono-S (Cytiva).

Isopeptide-linked ubiquitinated H3 for cryo-EM single particle analysis was prepared using mouse UBA1 (E1), human UBE2D3 (E2), human UHRF1 (E3), ubiquitin and H3_1-37W_ harboring K14R/K27R/K36R mutations (H3_1-37W_ K14R/K27R/K36R), which were purified in house as previous report ^24^. The ubiquitination reaction mixture contained 0.2 μM E1, 0.8 μM E2, 3 μM E3, 150 μM ubiquitin, and 50 μM H3_1-37W_ K14R/K27R/K36R mutant in 1 mL of ubiquitination reaction buffer (50 mM Tris-HCl [pH 8.5], 50 mM NaCl, 5 mM MgCl_2_, 0.1% Triton X-100, and 2 mM DTT). The reaction mixture was incubated at 30 °C for 6 h, and thereafter the reaction was quenched by the heat shock at 70 °C for 30 min, followed by collecting soluble fraction (Extended Data Fig. 1a). 6 μM of 18 bp hemimethylated DNA was added in initially ubiquitination reaction to enhance the ubiquitination reaction, however, the DNA addition was omitted for the sample preparation of DNMT1:H3Ub2:DNA^mCG/fCG^ to prevent DNA contamination.

### Preparation of binary and ternary complex for cryo-EM single particle analysis

For preparation of DNMT1:H3Ub2 binary complex, 20 μM isopeptide bound-ubiquitinated H3_1-37_ mixed with 6.6 μM apo-DNMT1 was subjected to size excursion chromatography (Superdex^®^ 200 Increase 10/300 GL, Cytiva) equilibrated with the buffer (20 mM Tris-HCl [pH 7.5], 250 mM NaCl, 10 μM Zn(OAc)_2_, and 0.5 mM DTT).

For preparation of ternary complex, purified DNMT1:H3Ub2 was mixed with 12 base pair of hemimetylated DNA analog (upper: 5’-ACTTA(5mC)GGAAGG, lower: 5’-CCTTC(5fC)GTAAGT: DNA^mCG/fCG^) in the conjugation buffer (50 mM Tris-HCl [pH 7.5], 20% Glycerol, 5 mM DTT and 50 mM NaCl). The conjugation reaction was initiated by the adding of 500 μM S-adenosyl-L-methionine (SAM) at 25 °C for 4 hrs. The yield was purified by size exclusion chromatography (Superdex^®^ 200 Increase 10/300 GL, Cytiva) equilibrated with cryo-EM buffer (20 mM Tris-HCl [pH 7.5], 250 mM NaCl and 5 mM DTT).

### Cryo-EM data collection

A 3 µL of the protein solutions was applied onto the glow-discharged holey carbon grids (Quantifoil Cu R1.2/1.3, 300 mesh). The grids were plunge-frozen in liquid ethane using a Vitrobot Mark IV (Thermo Fisher Scientific). Parameters for plunge-freezing were set as follows: blotting time, 3 sec; waiting time, 0 sec; blotting force, −0; humidity, 100%; and chamber temperature, 4 °C. Data for DNMT1:H3Ub2:DNA^mCG/fCG^ ternary complex was collected at the University of Tokyo on a 300 kV Titan Krios electron microscope (Thermo Fisher Scientific) with a K3 direct electron detector (Gatan) with BioQuantum energy filter in counting mode. A total of 4,068 movies were recorded at nominal magnification of ×105,000 and a pixel size of 0.83 Å/pixel, with a defocus range between −0.8 and −1.8 μm and a dose rate of 1.25 electrons/Å^2^ per frame. A typical motion-corrected cryo-EM image is shown in Supplementary Fig. 1a.

Data of apo-DNMT1, DNMT1:H3Ub2 were collected at RIKEN BDR on a 200-kV Tecnai Arctica electron microscope (Thermo Fisher Scientific) with a K2 direct electron detector (Gatan) in counting mode. A total 2,071 movies for apo-DNMT1 and 1,869 movies for DNMT1:H3Ub2 were recorded at nominal magnification of ×23,500 and a pixel size of 1.477 Å/pixel, with a defocus range between −0.8 and −1.4 μm, and a dose rate of 1.25 electrons/Å^2^ per frame. A typical motion-corrected cryo-EM image is shown in Supplementary Figs. 2a and 3b.

### Data processing

All data were processed using cryoSPARC v3.2.0 ^29^ for PDB deposition. The movie stacks were motion corrected by Full-frame motion correction or Patch motion correction. The defocus values were estimated from Contrast transfer function (CTF) by Patch CTF estimation or CTFFIND4 ^30^. A total of 4,307,107 particles of DNMT1:H3Ub2:DNA^mCG/fCG^ ternary complex were automatically picked using blob picker with 80 Å, 105 Å and 130 Å circular blobs and 80-130 Å elliptical blobs (Supplementary Fig. 1b). Particles (3,798,046) were then extracted in a box size of 256 pixels with a 0.83 Å/pixel size followed by single round of reference-free 2D classifications (Supplementary Fig. 1c). The selected good class containing 1,621,988 particles were used for *ab initio* 3D reconstruction (Extended Date Fig. 3). Then, non-uniform refinement was performed against all the extracted particles to yield the cryo-EM map with an overall resolution of 2.09 Å resolution. The subsequent heterogeneous refinement selected 2,653,627 particles as a good class. These particles were subjected to a 3D variability analysis, separating the CXXX-ordered and CXXC-disordered models. The particles (138,662) in the CXXC-ordered model and those (897,446) in the CXXC-disordered models were then subjected to non-uniform refinement to generate a cryo-EM map with an overall resolution of 2.52 Å and 2.23 Å, respectively. The classification processes were shown in Extended Data Fig. 3, and the statics of data collection and refinement and validation were shown in Table 1.

A total of 3,984,637 particles of apo-DNMT1 were automatically picked using blob picker with 80 Å, 105 Å and 130 Å circular blobs (Supplementary Fig. 2b). Particles (3,984,637) were then extracted in a box size of 160 pixels with a 1.477 Å/pixel size, followed by two rounds of reference-free 2D classifications to remove junk particles (Supplementary Fig. 2c). The selected 3,666,067 particles were subjected for *ab initio* 3D model reconstruction to generate four cryo-EM maps (Supplementary Fig. 4). An initial model shows similar shape with the crystal structure of DNMT1 (PDB: 4WXX). Then, non-uniform refinement was performed against the particles classified in the initial model of DNMT1 (1,824,727). These particles were re-extracted in a box size of 256 pixels with a 1.477 Å/pixel size by local motion correction. Non-uniform refinement yields the cryo-EM map with an overall resolution of 3.32 Å resolution. These particles were subjected to a 3D variability analysis and heterogeneous refinement, removing the dimer particles, and separating RFTS-free map and RFTS-bound map. To improve the cryo-EM map, further 3D variability analysis and clustering by PCA was performed. The particles (380,989) were then subjected to non-uniform refinement to yield final cryo-EM map. Overall resolution of 3.45 Å resolution using the gold-standard Fourier shell correlation with a 0.143 cut-off. The classification processes were shown in Supplementary Fig. 4, and the statics of data collection and refinement and validation were shown in Supplementary Table 1.

A total of 2,463,410 particles of DNMT1:H3Ub2 were automatically picked using blob picker with 80 Å, 105 Å and 130 Å circular blobs (Supplementary Fig. 3b). The particles were then extracted in a box size of 160 pixels in a 1.477 Å/pixel size using cryoSPARC followed by initial dataset cleanup using reference-free 2D classifications (Supplementary Fig. 3c). The selected 2,336,267 particles were subjected for *ab initio* 3D model reconstruction to generate four cryo-EM maps (Supplementary Fig. 5). Further 2D classifications were performed by the 2 class of *ab initio* 3D model assigned as DNMT1 particles. The particles (160,088) belong with the best four 2D classes were subjected to create the fine initial model. For the further refinement, the particles (1,303,645) without ice images from 2D classification II were selected. These particles were re-extracted in a box size of 256 pixels with a 1.477 Å/pixel size by local motion correction. Non-uniform refinement yields the cryo-EM map with an overall resolution of 3.55 Å resolution. These particles were subjected to a 3D variability analysis and heterogeneous refinement, separating RFTS-free map and RFTS-bound map. The subsequent 2D classification selected 735,233 particles as RFTS-bound structure. To improve the cryo-EM map, further 3D variability analysis and clustering by PCA was performed. The particles (645,368) were then subjected to non-uniform refinement to yield final cryo-EM map. Overall resolution of 3.52 Å resolution using the gold-standard Fourier shell correlation with a 0.143 cut-off. The classification processes were shown in Supplementary Fig. 5, and the statics of data collection and refinement and validation were shown in Supplementary Table 1.

For the analysis of H3Ub2-RFTS domain complex, the 2D classification analysis were also performed by Relion 3.1 (Extended Data Fig. 6) ^31^. The movie stacks of DNMT1:H3Ub2:DNA^mCG/fCG^ were motion corrected by MotionCor2. The defocus values were estimated from Contrast transfer function (CTF) by CTFFIND4 ^30^.Single particle image was also extracted by LoG Auto picker of Relion to check the particles of other biomolecules. After three rounds of 2D classification, the particle smaller than DNMT1 was selected. These smaller particles (80,186) were re-extracted in a box size of 128 pixels with a 0.83 Å/pixel size, and the images were classified by 2D classification. The major 2D average images were compared with the projected templates of RFTS-H3Ub2 complex (PDB: 5WVO) (Extended Data Fig. 6). Gaussian model of the complex was created by the Molmap of ChimeraX ^32^. The 2D projected templates were created by the module of “create template” in cryoSPARC.

### SEC-SAXS

SAXS data were collected on Photon Factory BL-10C using a HPLC Nexera/Prominence-I (Shimazu) integrated SAXS set-up. 100 µL of a 10 mg/mL of the apo-DNMT1 (aa:351-1616) or its bound to H3Ub2^S-S^ were loaded onto a Superdex^®^ 200 Increase 10/300 GL (Cytive) pre-equilibrated with 20 mM Tris-HCl (pH 8.0), 150 mM NaCl and 5% glycerol at a flow rate of 0.5 mL/min at 20 °C. The flow rate was reduced to 0.05 mL/min at an elution volume of 10 −13 mL. X-ray scattering was collected every 20 seconds on a PILATUS3 2M detector over an angular range of *q*_min_ = 0.00690 Å^−1^ to *q*_max_ = 0.27815 Å^−1^. UV spectra at a range of 200 nm to 450 nm were recorded every 10 seconds. Circular averaging and buffer subtraction were carried out using the program SAngler ^33^ to obtain one-dimensional scattering data *I*(*q*) as a function of *q* (*q* = 4πsinθ/λ, where 2θ is the scattering angle and λ is the X-ray wavelength 1.5 Å). The scattering intensity was normalized on an absolute scale using the scattering intensity of water ^34^. The multiple concentrations of the scattering data around the peak at A280, namely ascending and descending parts of chromatography peak, and *I*(0) were extrapolated to zero-concentration by Serial Analyzer ^35^. The molecular mass of the measured proteins was estimated by the empirical volume of correlation, *V*_c_, showing no aggregation of the measured sample ^36^. The radius of gyration *R*_g_ and the forward scattering intensity *I*(0) were estimated from the Guinier plot of *I*(*q*) in the smaller angle region of *qR*_g_ < 1.3. The distance distribution function *P*(*r*) was calculated using the program GNOM ^37^. The maximum particle dimension *D*_max_ was estimated from the *P*(*r*) function as the distance *r* for which *P*(*r*) = 0. The scattering profile of crystal structure of apo-DNMT1 and its docking model with ubiquitinated H3 were computed with CRYSOL ^38^.

### *In vitro* DNA methylation assay

The 42 base pair of DNA duplex containing three hemimethylation sites (0 - 0.8 µM) was methylated with the recombinant DNMT1 (15 nM, aa:351-1616) by the addition of the disulfide-linked ubiquitinated H3 (1 µM H3Ub2^S-S^) including 20 µM SAM in reaction buffer (20 mM Tris-HCl [pH 8.0], 50 mM NaCl, 1 mM EDTA, 3 mM MgCl_2_, 0.1 mg/mL BSA and 20% Glycerol) at 37 °C for 1 hr. Termination of methylation reaction and conversion of SAH to ADP were performed by the addition of 5×MTase-Glo^TM^ reagent from methyltransferase assay kit, MTase-Glo (Promega) at 1:4 ratio for the reaction total volume. After 30 min stationary at room temperature, ADP detection process was carried out with solid white flat bottom 96 well plates (Costar). MTase-Glo^TM^ Detection Solution was added to the reaction in 1:1 ratio to reaction total 40 µL volume and incubated for 30 min at room temperature. The luminescence derived from reaction product, SAH, was monitored using GloMax® Navigator Microplate Luminometer (Promega). The effect of F631A/F632A mutation were examined at the condition of the DNMT1 (15 nM) with 1 µM H3Ub2^S-S^ by the addition of 42 base pair of DNA duplex (0 - 0.8 µM) in same reaction buffer. The SAH conversion process and ADP detection process in the manner described above.

For the evaluation of the DNMT1:H3Ub2 complex, final concentration of DNMT1 was 50 nM to prevent the dissociation of ubiquitinated H3 (*K*_D_ = 18 nM). DNA methylation reactions were initiated by mixing of apo-DNMT1 or DNMT1:H3Ub2^iso^ and the stopped at 0, 5, 15 or 30 min by addition of 5×MTase-Glo^TM^ reagent. The detection process was performed in the same way as described above. At least three independent experiments were performed for estimation of standard deviation.

### DNA pull-down assay

20 µg of the 21 base pair of biotinylated hemimethylated DNA duplex was immobilized on Dynabeads M-280 Streptavidin (VERITAS) equilibrated with the binding buffer (20 mM Tris-HCl [pH 7.5], 150 mM NaCl, 10% Glycerol and 0.05% Nonidet P-40 (NP-40)). After washing the beads with the binding buffer, 10 µg of purified DNMT1 (aa:351-1616) wild-type or F631A/F632A mutant, 2-equimolar excess of H3Ub2^S-S^ and equimolar of SAH were added to the beads. After incubation for 2 hr at 4 °C, the unbound proteins were washed five times with the binding buffer. The proteins bound to the immobilized DNA were boiled for 2 min at 95 °C in an oxidative SDS-loading buffer and analyzed by SDS-PAGE using SuperSep^TM^ Ace, 5-20% gel (Wako, Japan). At least three independent experiments were performed.

### *Xenopus* egg extracts

*Xenopus laevis* was purchased from Kato-S Kagaku and handled according to the animal care regulations at the University of Tokyo. The preparation of interphase egg extracts, chromatin isolations, and immunodepletions were performed as described previously ^24, 39^. Unfertilized *Xenopus laevis* eggs were dejellied in 2.5% thioglycolic acid-NaOH (pH 8.2) and washed in 1xMMR buffer (100 mM NaCl, 2 mM KCl, 1 mM MgCl_2_, 2 mM CaCl_2_, 0.1 mM EDTA, 5 mM HEPES-NaOH [pH 7.5]). After activation in 1xMMR supplemented with 0.3 µg/mL calcium ionophore, eggs were washed with EB buffer (50 mM KCl, 2.5 mM MgCl_2_, 10 mM HEPES-KOH [pH 7.5], 50 mM sucrose). Eggs were packed into tubes by centrifugation (BECKMAN, Avanti J-E, JS-13.1 swinging rotor) for 1 min at 190×g and crushed by centrifugation for 20 min at 18,973×g. Egg extracts were supplemented with 50 µg/mL cycloheximide, 20 µg/mL cytochalasin B, 1 mM DTT, 2 µg/mL aprotinin, 5 µg/mL leupeptin and clarified by ultracentrifugation (Hitachi, CP100NX, P55ST2 swinging rotor) for 20 min at 48,400×g. The cytoplasmic extracts were aliquoted, frozen in liquid nitrogen, and stored at −80 °C. All extracts were supplemented with an energy regeneration system (2 mM ATP, 20 mM phosphocreatine, and 5 µg/ml creatine phosphokinase). 3,000-4,000 nuclei/µl of sperm nuclei were added and incubated at 22 °C. Aliquots (15-20 µl) were diluted with 150 µl chromatin purification buffer (CPB; 50 mM KCl, 5 mM MgCl_2_, 20 mM HEPES-KOH [pH 7.6]) containing 0.1% NP-40, 2% sucrose, 2 mM N-ethylmaleimide (NEM). After incubation on ice for 5 min, diluted extracts were layered over 1.5 mL of CPB containing 30% sucrose and centrifuged at 15,000×g for 10 min at 4 °C. Chromatin pellets were resuspended in 1×Laemmli sample buffer, boiled for 5 min at 100 °C, and analyzed by immunoblotting.

For protein expression in insect cells, baculoviruses were produced using a BD BaculoGold Transfection kit and a BestBac Transfection kit (BD Biosciences), following the manufacturer’s protocol. Proteins were expressed in Sf9 insect cells by infection with viruses expressing xDnmt1 Wt-FLAGx3, xDnmt1 F506AF507A-FLAGx3 for 72 hrs. Sf9 cells from a 500 ml culture were collected and lysed by resuspending in 20 mL lysis buffer (20 mM Tris-HCl [pH 8.0], 100 mM KCl, 5 mM MgCl_2_, 10% glycerol, 1% NP-40, 1 mM DTT, 10 µg/mL leupeptin, and 10 µg/mL aprotinin), followed by incubation on ice for 10 min. A soluble fraction was obtained after centrifugation of the lysate at 15,000×g for 15 min at 4 °C. The soluble fraction was incubated for 4 hrs at 4 °C with 250 µL of anti-FLAG M2 affinity resin (Sigma-Aldrich) equilibrated with lysis buffer. The beads were collected and washed with 10 mL wash buffer (20 mM Tris-HCl [pH 8.0], 100 mM KCl, 5 mM MgCl_2_, 10% glycerol, 0.1% NP-40, 1 mM DTT) and then with 5 mL EB (20 mM HEPES-KOH [pH 7.5], 100 mM KCl, 5 mM MgCl_2_) containing 1 mM DTT. The recombinant xDnmt1 was eluted twice in 250 µL EB containing 1 mM DTT and 250 µg/mL 3×FLAG peptide (Sigma-Aldrich). Eluates were pooled and concentrated using a Vivaspin 500 (GE Healthcare Biosciences).

### Cell Culture, Transfection, and Colony Isolation

The HCT116 cell line, which conditionally expressed OsTIR1 under the control of a tetracycline (Tet)-inducible promoter, was obtained from the RIKEN BRC Cell Bank (http://cell.brc.riken.jp/en/), and genotyped by Eurofins. All cell lines were cultured in McCoy’s 5A medium (Sigma-Aldrich) supplemented with 10% FBS (Gibco), 2 mM L-glutamine, 100 U/mL penicillin, and 100 µg/mL streptomycin. Cells were grown in a 37 °C humid incubator with 5% CO_2_. To generate stable DNMT1-AID cell lines, we followed previous studies ^40, 41^. Briefly, cells were grown in a 24 well plate, then CRISPR/Cas and donor plasmids were transfected using Lipofectamine 2000 (Thermo Fisher Scientific). Two days after transfection, cells were transferred and diluted in 10 cm dishes, followed by selection in the presence of 700 mg/mL G418 or 100 mg/mL Hygromycin B. After 10–12 days, colonies were picked for further selection in a 96-well plate. To induce the degradation of AID-fused proteins, cells were incubated with 0.2 µg/mL doxycycline (Dox) and 20 µM auxinole for 1 day, then we replaced the medium including 0.2 µg/mL Dox and 500 µM indole-3-acetic acid (IAA), a natural auxin.

### Establishment of stable expressing exogenous DNMT1 cell lines for rescue experiments

We cloned WT DNMT1 and point mutant DNMT1 (F631A/F632A) to pSBbi-Bla (Addgene: 60526). All plasmids were sequenced prior to use. To establish stable expressing exogenous DNMT1 cell lines, we used sleeping beauty system^42^. Briefly, cells were grown in a 6 well plate, then transposase vector (Addgene: 34879) and each pSBbi-Bla plasmids (EV, WT, F631A/F632A) were transfected using Lipofectamine 2000 (Thermo Fisher Scientific). Four days after transfection, cells were selected with 10 µg/mL Blasticidin for one week. To detect DNA methylation level, LUMA and Pyrosequencing were done according to standard procedures.

### Data availability

The cryo-EM density map has been deposited in the Electron Microscopy Data Bank (EMDB, www.ebi.ac.uk/pdbe/emdb/) with the accession code EMD-33200, EMD-33201, EMD-33298, EMD-33299 and the atomic coordinates have been deposited in the PDB (www.rcsb.org) with the accession code 7XI9, 7XIB. All data needed to evaluate the conclusions in the paper are present in the paper and/or the Supplementary Materials. Additional data related to this paper may be requested from the authors.

## Acknowledgments

The cryo-EM experiments were performed at the cryo-EM facility of the RIKEN Center for Biosystems Dynamics Research Yokohama. We thank the members of Structure Biology Research Center at KEK, especially H. Kawasaki and T. Senda for Cryo-EM and N. Shimizu for SAXS. We thank the members of cryo-EM facility at the University of Tokyo, especially Y. Sakamaki and M. Kikkawa. M. Ariyoshi supports in the preparation of the manuscript.

This research was partially supported by Platform Project for Supporting Drug Discovery and Life Science Research (Basis for Supporting Innovative Drug Discovery and Life Science Research [BINDS]) from AMED under Grant Number 1770, 2102, 2965 and 3002. PAD is supported by Agence Nationale de la Recherche (PRCI INTEGER ANR-19-CE12-0030-01), LabEx “Who Am I?” (ANR-11-LABX-0071), Université de Paris IdEx (ANR-18-IDEX-0001) funded by the French Government through its “Investments for the Future” program, Fondation pour la Recherche Médicale, Fondation ARC (Programme Labellisé PGA1/RF20180206807). KY was the recipient of a postdoctoral fellowship from Fondation Association pour la Recherche sur le Cancer, and of a subsequent postdoc fellowship from Labex “Who Am I?”.

## Author information

These authors contributed equally: Amika Kikuchi, Hiroki Onoda,

## Contributions

K.A. supervised the work. A.K., H.O., S.K., S.M., S.Y., and H.S. performed cloning and protein purification. A.K., H.O., S.Y., H.S. performed *in vitro* biochemical assay. S.K., and K.A. performed SEC-SAXS experiments and analysis. A.K., H.O., A.Y., M.S., N.A., K.A. performed Cryo-EM experiments and analysis. Y.C., S.T., M.N., and A.N. performed evaluation of mutant using *Xenopus* egg extract and K.Y., J.S., H.K., and PAD performed cell-based assay. H.O., PAD, and K.A. wrote the paper.

## Corresponding author

Correspondence to Kyohei Arita

## Competing interests

Authors declare that they have no competing interests.

---

**Extended Data Fig. 1.**
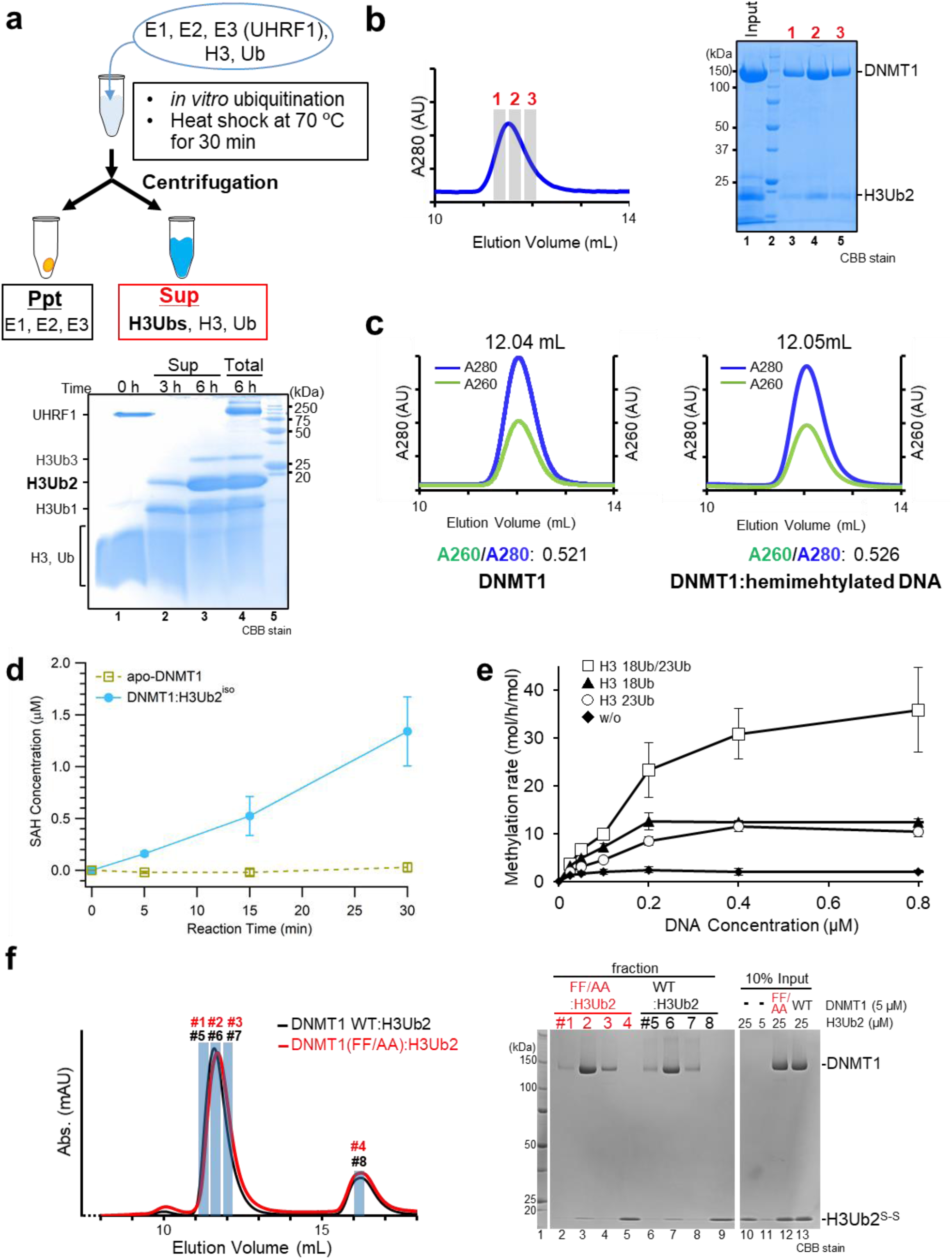
Sample preparation of DNMT1:H3Ub2 and biochemical assay. **(a)** Schematic figure of *in vitro* preparation of ubiquitinated H3. Lower panel shows analysis of ubiquitination of histone H3_1-37W_ by SDS-PAGE. Lane 1: ATP omitted-reaction cocktail, lanes 2 and 3: Supernatant (sup) of *in vitro* ubiquitination reaction cocktail for 3 h and 6 h after heat-shock at 70 °C, lanes 4: Total proteins in the reaction cocktail for 6 h reaction before the heat shock, lane 5: size marker. **(b)** Size exclusion chromatography of the mixture of DNMT1_351-1616_ and excess ubiquitinated H3. Vertical and horizontal axes indicate the absorbance of 280 nm (blue line: A280) and elution volume of Superdex^®^ 200 Increase 10/300 GL, respectively. SDS-PAGE of input (lane 1) and elution fractions were depicted in the chart of size exclusion chromatography. **(c)** Size exclusion chromatography of apo-DNMT1 (left) and apo-DNMT1 mixed with hemimethylated ^mCG/fCG^ DNA (DNA) after conjugating reaction for 12 hr at 25 °C (right). A260/A280 value at the peak is shown, indicating that DNMT1 was unable to form the binary complex with the DNA. **(d)** DNA methylation assay of DNMT1 (aa:351-1616) in the absence of and presence of H3Ub2^iso^. Vertical and horizontal axes indicate SAH concentration and reaction time, respectively. Line shows the average value of three times independent experiments. **(e)** DNA methylation assay of apo-DNMT1 (black-square) and DNMT1 in the presence of H3K18Ub (black-triangle), H3K23Ub (white-circle) and H3K18Ub/K23Ub (white-square). At least three independent experiments were performed for estimation of standard deviation. **(f)** Left: Size exclusion chromatography of the mixture of wild type (WT) or mutant (FF/AA: F631A/F632A) DNMT1_351-1616_ in the presence of excess H3Ub2^S-S^. Vertical and horizontal axes indicate the absorbances at 280 nm and elution volume from Superdex^®^ 200 Increase 10/300 GL, respectively. Right: Analysis of complex formation of WT or FF/AA mutant DNMT1 with H3Ub2^S-S^ by SDS-PAGE. Lanes 10-13 show the input sample for the SEC analysis.

**Extended Data Fig. 2.**
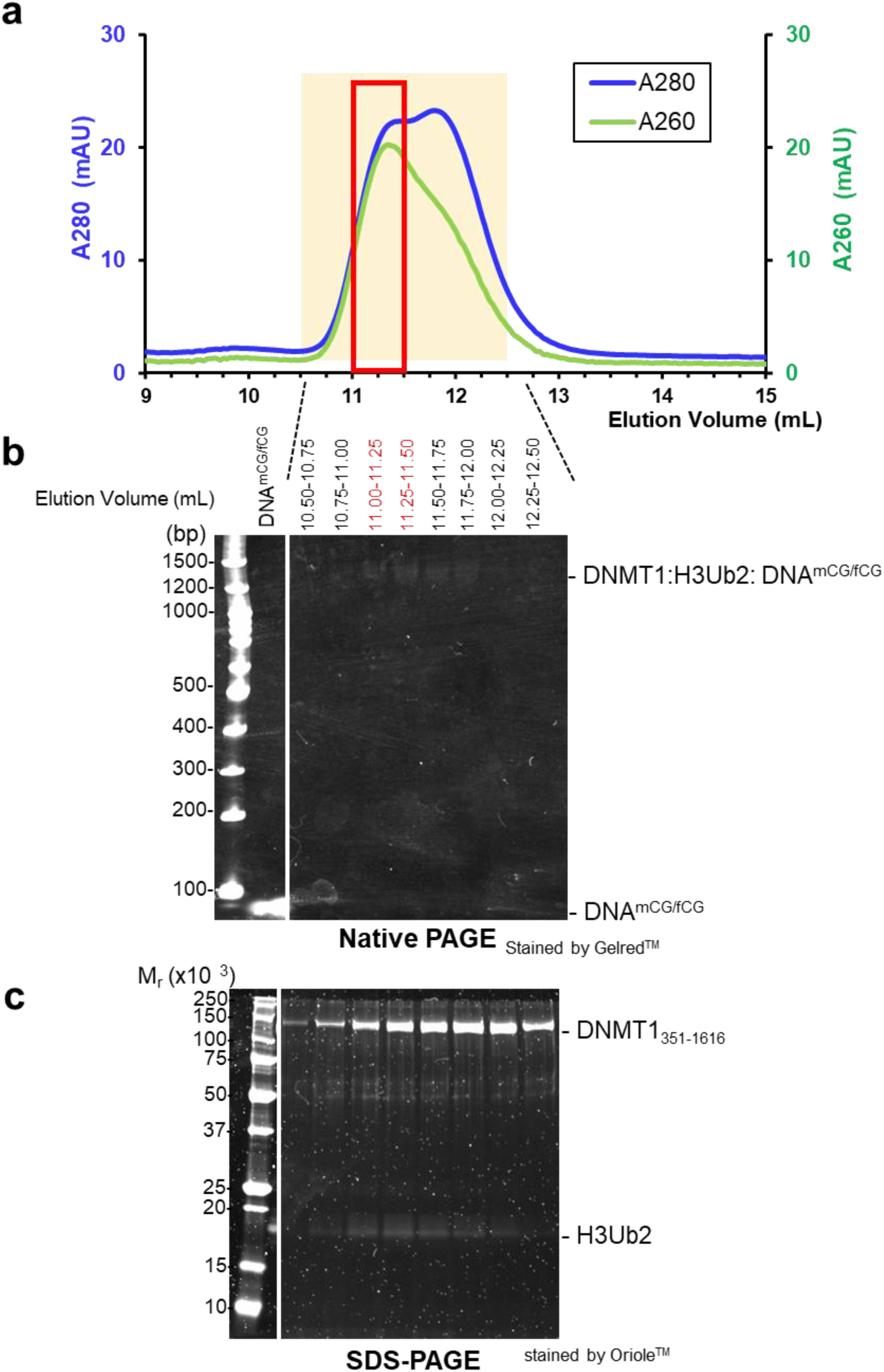
Sample preparation of DNMT1:H3Ub2:DNA^mCG/fCG^ ternary complex for cryo-EM. **(a)** Size exclusion chromatography of the mixture of DNMT1:H3Ub2 binary complex and DNA^mCG/fCG^. All fractions in the chromatography peaks were analyzed by **(b)** native-PAGE stained by Gelred^TM^ for detecting DNA and **(c)** SDS-PAGE stained by Oriole^TM^ for detecting proteins.

**Extended Data Fig. 3.**
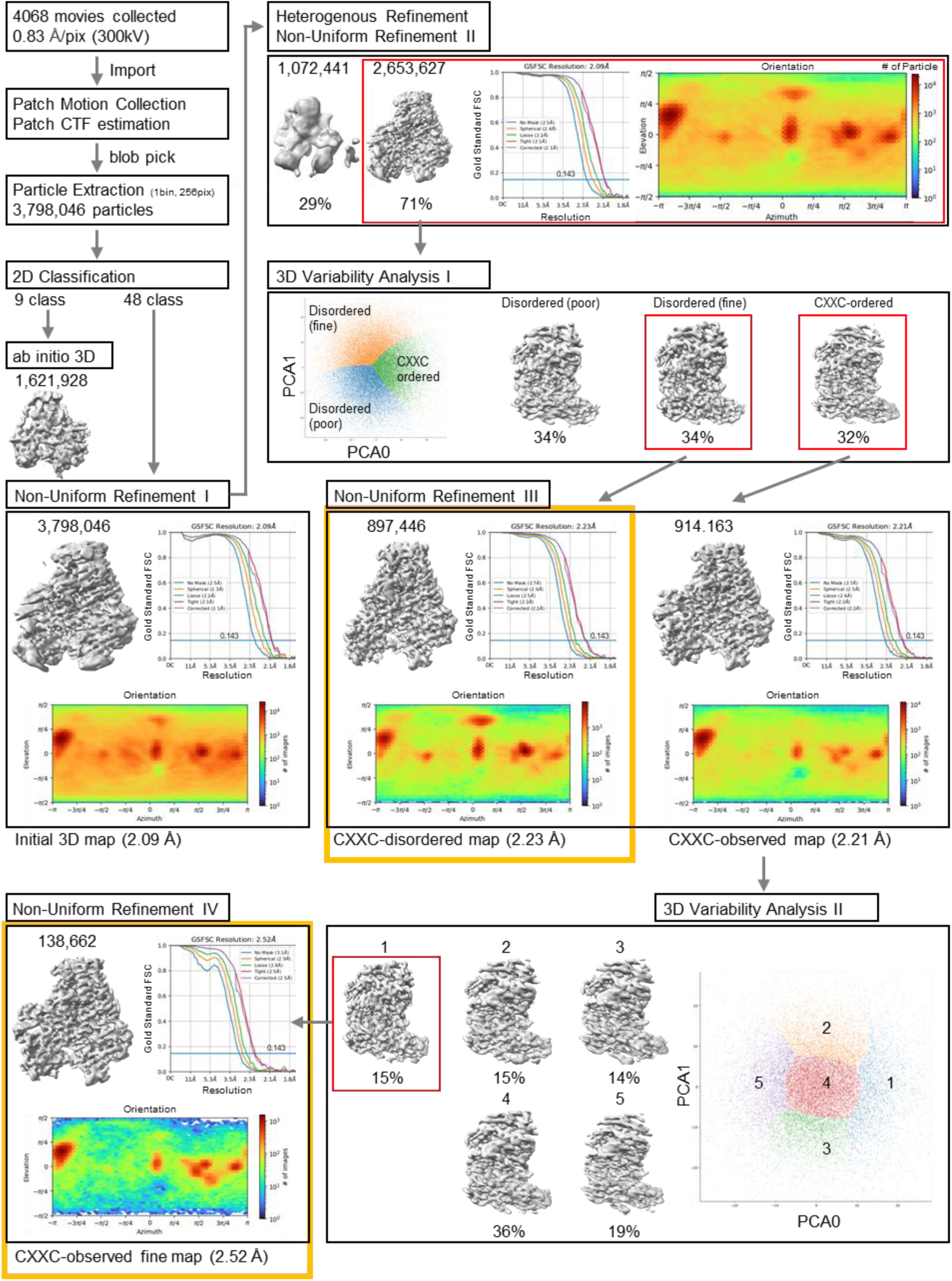
Overview of cryo-EM data processing workflow for DNMT1:H3Ub2:DNA^mCG/fCG^. Data processing were performed on cryoSPARC. 3,798,046 particles were automatically picked blob picker from 4,068 motion corrected movies. After the reference-free 2D classification, *ab initio* 3D model was constructed from the best nine in the 2D classes. Then, Non-Uniform Refinement I was performed against all the extracted particles to yield the cryo-EM map with an overall resolution of 2.09 Å resolution. The subsequent heterogeneous refinement selected 2,653,627 particles as a good class. These particles were subjected to a 3D variability analysis, separating the CXXC-ordered and CXXC-disordered models. The particles (138,662) in the CXXC-ordered model and those (897,446) in the CXXC-disordered models were then subjected to Non-Uniform Refinement (III, IV) to generate a cryo-EM map with an overall resolution of 2.52 Å and 2.23 Å, respectively.

**Extended Data Fig. 4.**
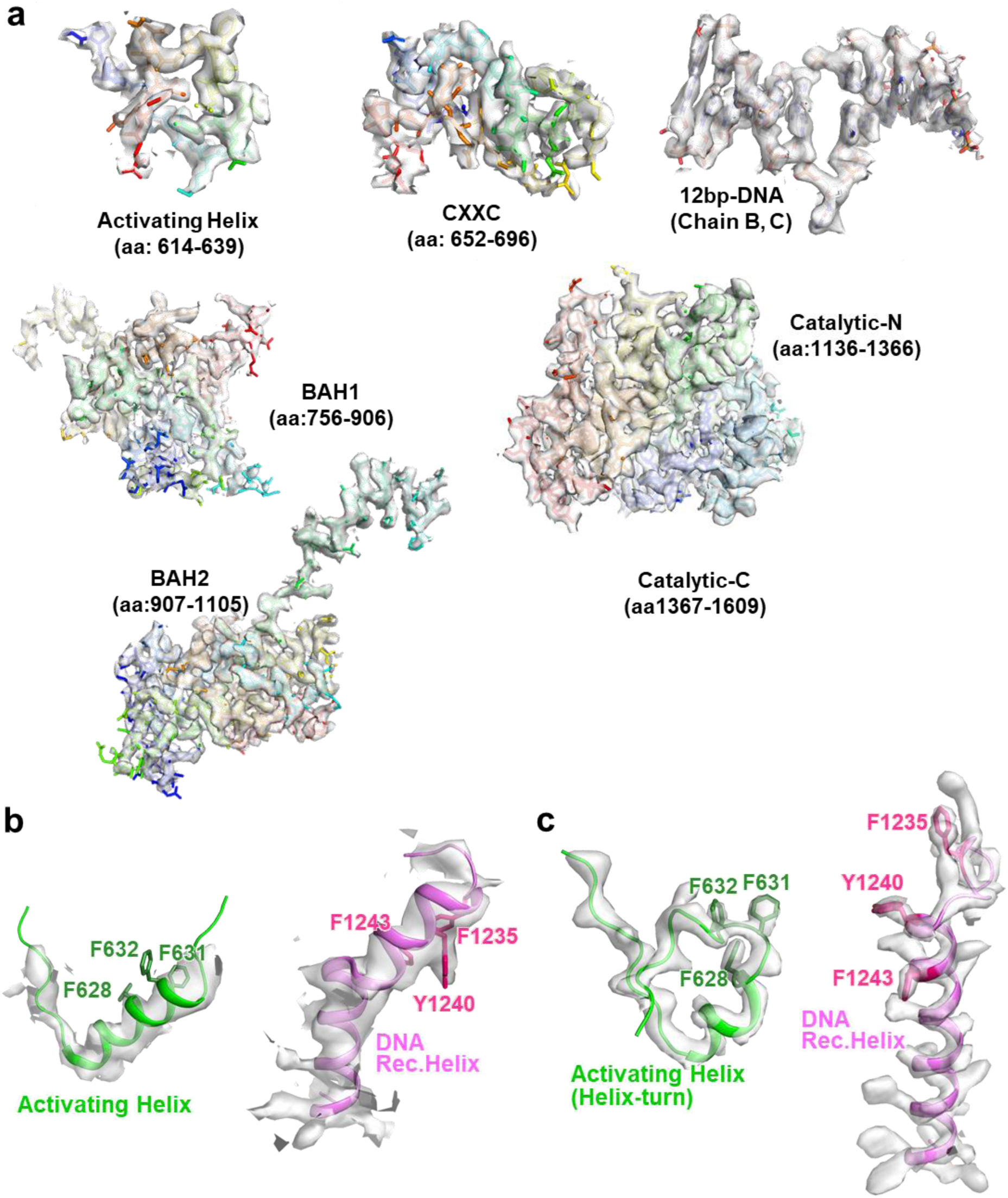
Cryo-EM maps of CXXC-ordered DNMT1:H3Ub2:DNA^mCG/fCG^. **(a)** The atomic models of DNMT1 and DNA are superimposed on the cryo-EM map. **(b)** Cryo-EM map of Activating Helix and DNA Recognition Helix of apo-DNMT1 (EMD-33299) superimposed on crystal structure of apo-DNMT1 (PDB:4WXX) **(c)** Structure of Activating Helix and DNA Recognition Helix in the ternary complex superposed on cryo-EM map (EMD-33200).

**Extended Data Fig. 5.**
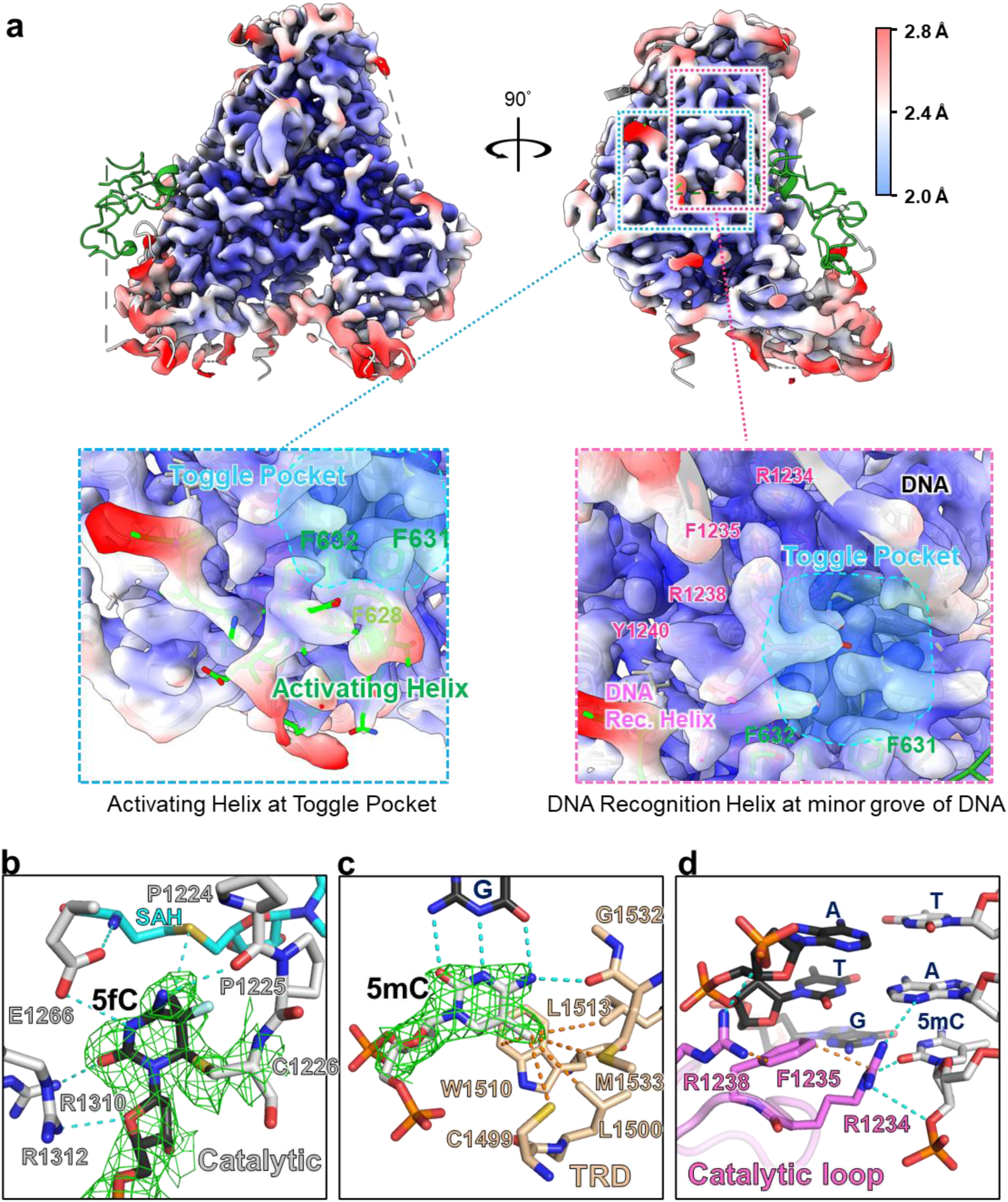
Local resolution map of CXXC-disordered DNMNT1:H3Ub2:DNA. **(a)** CXXC-disordered cryo-EM map of transparent surface were overlayed on the cartoon model of CXXC-ordered model by UCSF Chimera. CryoEM map from front view (upper left) and side view (upper right) were colored with the local resolution, which is estimated by cryoSPARC. Close-up view of the Toggle Pocket (lower left) and minor groove of DNA at mCG/fCG site (lower right) were depicted in lower panel. Stick model of CXXC-ordered model were shown in same color scheme with Fig. 1b. **(b)** Recognition of 5-fluolocytosine (5fC). 5fC, amino acid residues in the catalytic domain of DNMT1 and SAH are shown as black, gray and cyan stick models, respectively. Cyan dotted line indicates hydrogen bond. Green mesh shows cryo-EM map. **(c)** 5-methylcytosine (5mC) recognition by TRD domain of DNMT1 showing light orange stick model. Cryo-EM map corresponding to 5mC was shown as green mesh. **(d)** Recognition of the DNA from the minor groove at mCG/fCG site by catalytic loop of DNMT1. The residues in the catalytic loop are shown as pink stick model.

**Extended Data Fig. 6.**
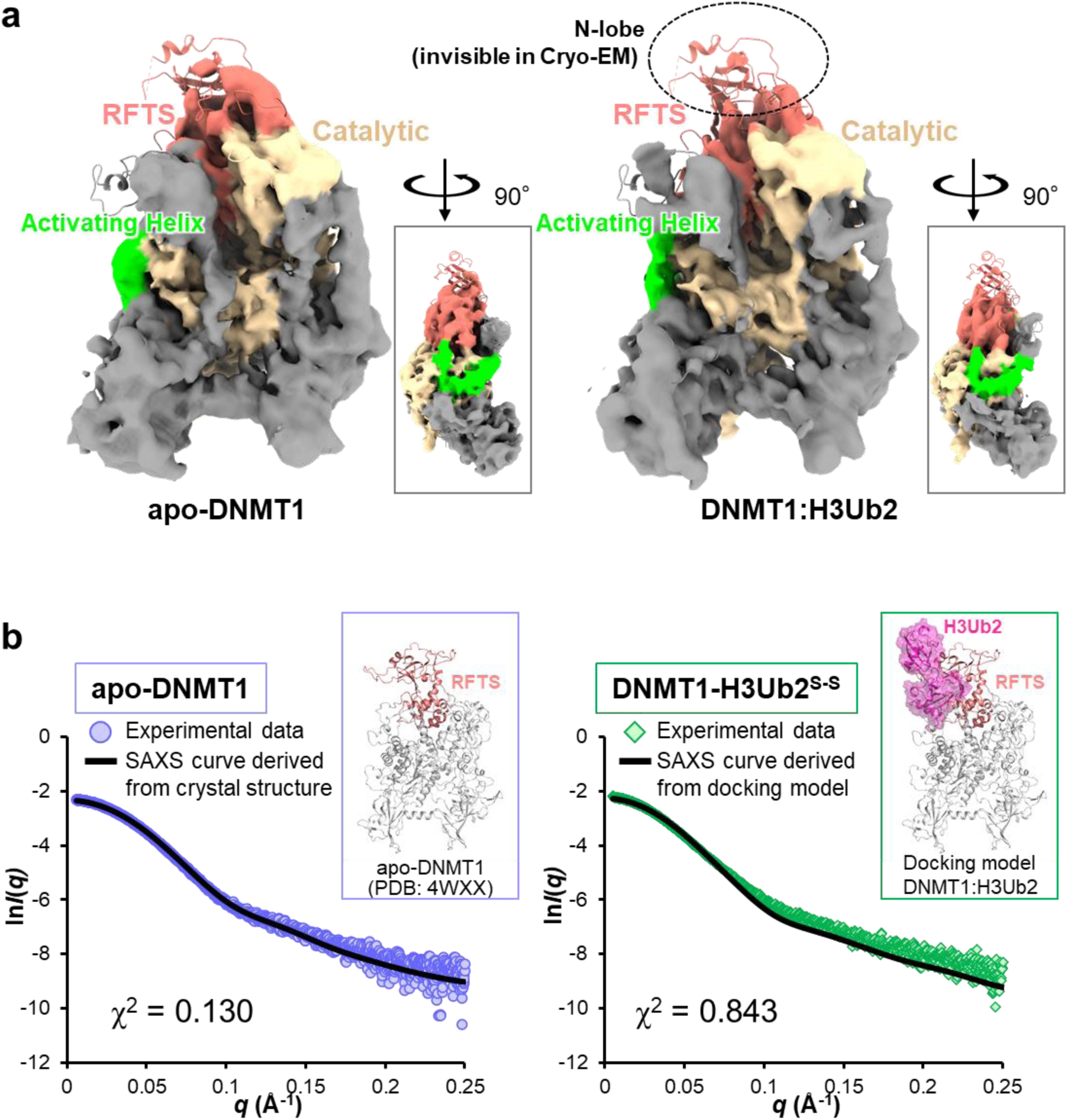
Structural analysis of apo-DNMT1 and it complex with H3Ub2. **(a)** Cryo-EM map of apo-DNMT1 (aa:351-1616, left) and the DNMT1:H3Ub2 binary complex (right). RFTS, Activating Helix and catalytic domains are colored salmon pink, green and beige, respectively. The cartoon model of crystal structure of apo-DNMT1 (PDB: 4WXX) is superimposed on the cryo-EM maps (EMD-33299, EMD-33298). **(b)** SAXS intensity data of apo-DNMT1 (purple circle) and DNMT1:H3Ub2^S-S^ complex (green square) superimposed on the theoretical scattering curves, showing black line, derived from crystal structure of apo-DNMT1 (PDB: 4WXX) and a model structure of apo-DNMT1 docking with RFTS:H3Ub2 complex (PDB:5WVO), respectively. The inset depicts the cartoon model of apo-DNMT1 and the docking model of DNMT1:H3Ub2 complex.

**Extended Data Fig. 7.**
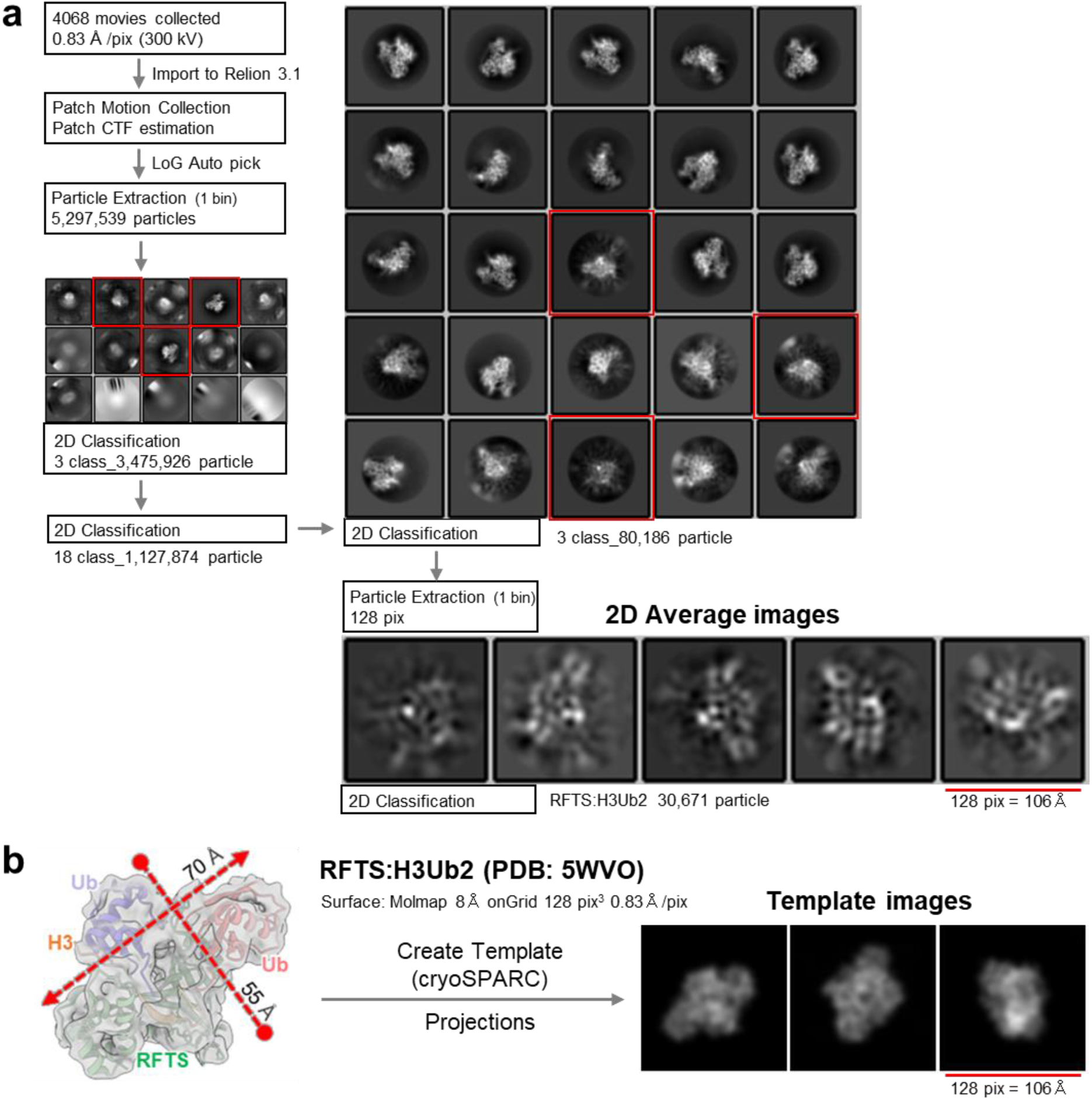
Cryo-EM analysis of DNMT1:H3Ub2:DNA^mCG/fCG^ using Relion 3.1. **(a)** Data processing were performed on Relion 3.1. Cryo-EM map refinement using Relion 3.1. 3,624,260 particles were automatically picked by crYOLO from 4,068 motion corrected movies. After the reference-free 2D classification, *ab initio* 3D model was constructed from the best nine in the 2D classes. The hand flipped *ab initio* 3D model of apo-DNMT1 (Supplemental Fig. 1) were used for the initial model. DNMT1 map was obtained after the single round of 2D classification and 2 rounds of Refine 3D. The final resolution was estimated as 2.3 Å by postprocess of Relion with soft mask. Single particle image was also extracted by LoG Auto picker of Relion to check the particles of other biomolecules. After three rounds of 2D classification, the particle smaller than DNMT1 was selected. These smaller particles (80,186) were re-extracted in a box size of 128 pixels with a 0.83 Å/pixel size, and the images were classified by 2D classification. **(b)** The major 2D average images were compared with the projected templates of RFTS:H3Ub2 complex (PDB: 5WVO). Gaussian model of the complex was created by the Molmap of ChimeraX. The 2D projected templates were created by the module of “create template” in cryoSPARC.

**Extended Data Fig. 8.**
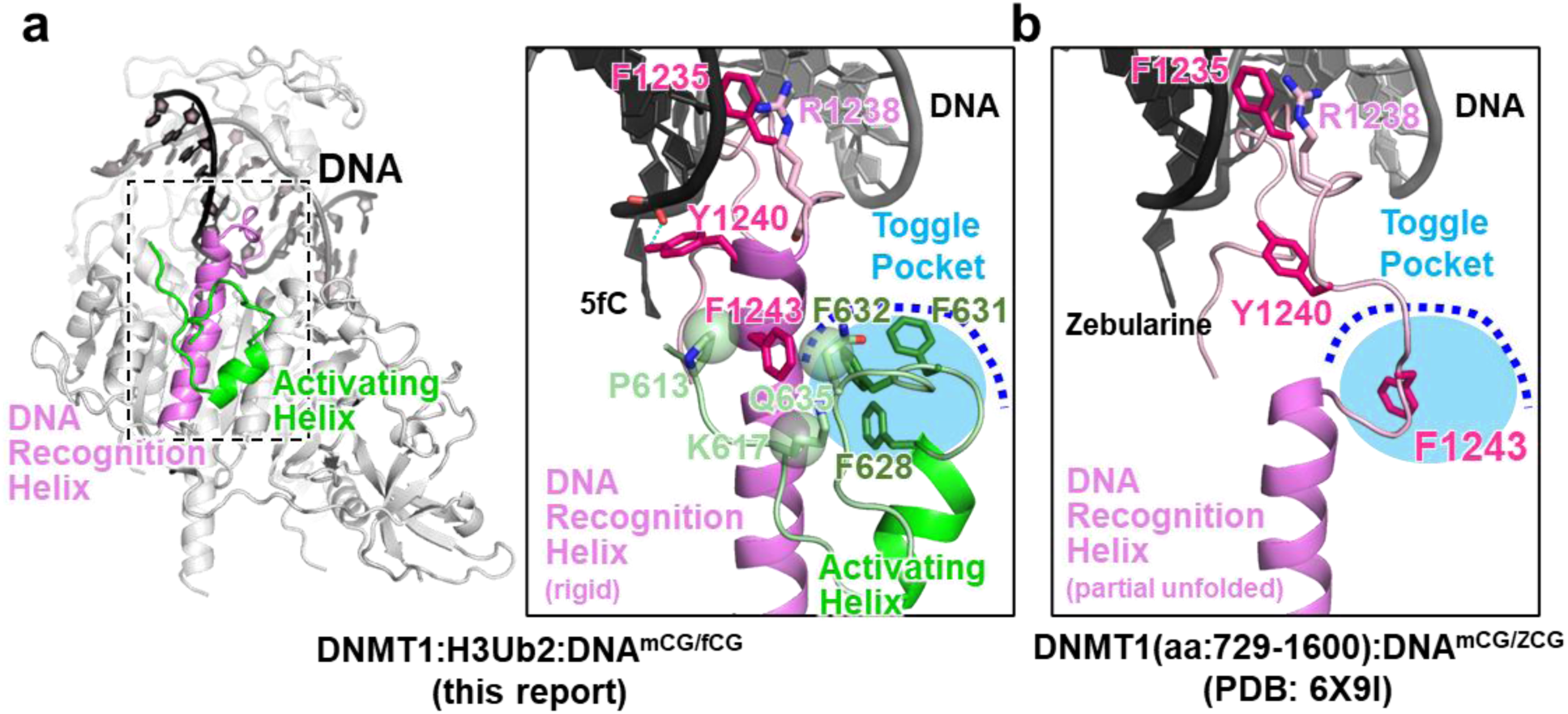
Structural comparison of DNA Recognition Helix in the ternary complex and DNMT1:DNAmCG/zCG binary complex (aa: 729-1600, PDB:6X9I). (a) Structure of DNA Recognition Helix (pink) and Activating Helix (green) of the ternary complex. Residues involving in recognition of DNA and interaction with the Toggle Pocket are shown as stick model. Toggle Pocket is highlighted as light blue. The cognate DNA strands are colored as black and gray. (b) Structure of DNA Recognition Helix of DNMT1:DNAmCG/zCG binary complex (z: Zebularine). Color schemes are same as (a).

**Extended Data Fig. 9.**
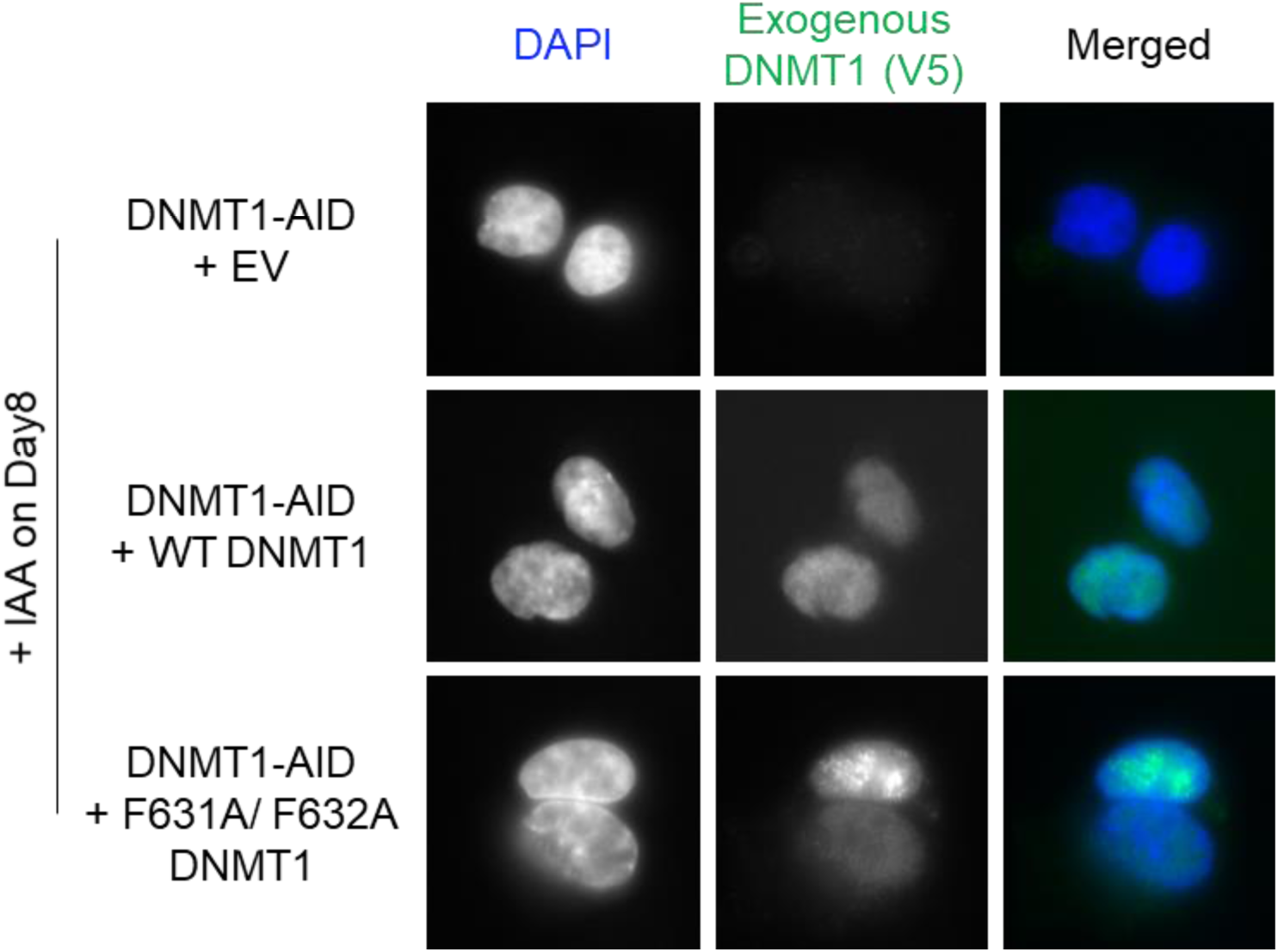
Nuclear localization of Exogenous DNMT1. The EV and exogenous V5 tagged DNMT1 (WT and F631A/F632A) were stably transfected to HCT116 cells expressing endogenous DNMT1-AID. The stable transfected cells were treated with IAA for 8 days. The nuclear was stained with DAPI, and exogenous DNMT1 was stained with V5 tag and second antibody. For merged images, the blue signal is corresponding to DAPI image, and the green signal is corresponding to exogenous DNMT1.

## Supplementary Notes

**Supplementary Note 1. Structural analysis of DNMT1 bound to double monoubiquitinated H3**

To examine the structural rearrangements of the DNMT1 protein upon binding of ubiquitinated H3 during enzymatic activation, we used small angle X-ray scattering (SAXS) and cryo-EM single particle analysis.

Histone H3 monoubiquitinated at K18 and K23 was prepared by linking the G76C ubiquitin by disulfide bond with H3 N-terminal tail harboring K18C/K23C (hereafter H3Ub2^s-s^) ^1^. We performed *in vitro* DNA methylation assay and found that binding of H3Ub2^s-s^ significantly enhanced the activity of DNMT1 (aa: 351-1616) (Extended Data Fig. 1e). Next, we conducted size exclusion chromatography in line with SAXS (SEC-SAXS) of apo-DNMT1 and DNMT1 bound to H3Ub2^s-s^ (Extended Data Fig. 6b, Supplementary Fig. 6, and Supplementary Table 2). The radius of gyration (*R*g) of apo-DNMT1 as estimated from Guinier analysis was 39.0 Å, and the scattering curve was well superimposed on the curve calculated from the crystal structure of human apo-DNMT1 (PDB: 4WXX). This indicated that in solution, apo-DNMT1 formed an autoinhibitory structure (Extended Data Fig. 6b left). The *R*g of the DNMT1 bound to H3-Ub2^s-s^ was modestly larger (43.7 Å) than that of the apo-DNMT1. However, the scattering curve of DNMT1 bound to H3Ub2^s-s^ was nearly identical to that of a model structure of DNMT1:H3Ub2 complex obtained by manual docking of RFTS:H3Ub2 structure (PDB: 5WVO) on the corresponding moiety of the crystal structure of apo-DNMT1. These results indicated that double monoubiquitinated H3 binding does not lead to displacement of the RFTS domain from the catalytic domain (Extended Data Fig. 6b right).

To further validate the above model, we performed cryo-EM single particle analysis of apo-DNMT1 bound to ubiquitinated H3. We first attempted preparation of the binary complex using DNMT1 and H3Ub2^S-S^; however, preparation of the cryogenic sample-grid suitable for atomic-resolution analysis failed under oxidative conditions. To overcome this problem, we performed *in vitro* ubiquitination and prepared isopeptide-linked ubiquitinated H3. H3 tail (residues 1-37W) containing the mutations K14R/K27R/K36R, prepared in house, was ubiquitinated using E1, E2 and UHRF1 (hereafter H3Ub2, see method). As expected, the binary complex DNMT1:H3Ub2 purified by a gel-filtration chromatography showed ubiquitinated H3-dependent DNA methylation activity (Extended Data Fig. 1d). Furthermore, 3.4 Å resolution cryo-EM map of apo-DNMT1 showed structural similarity with the crystal structure of apo-DNMT1 except for the flexible states of the CXXC domain and N-lobe of the RFTS domain (Extended Data Fig. 6a left). C-lobe of RFTS domain was embedded into the catalytic core, indicating an autoinhibitory state (Extended Data Fig. 6a left) ^2, 3^. 3.6 Å resolution of cryo-EM map of the DNMT1 in complex with H3Ub2 also revealed that the binding of H3Ub2 increased the dynamics of N-lobe of RFTS domain, resulting in complete invisibility of N-lobe bound to H3Ub2 (Extended Data Fig. 6a right). The density of N-lobe of the RFTS domain was obscured in the presence and absence of the H3Ub2 due to conformational flexibility of a long-α helix connecting the two lobes ^4^. However, the C-lobe was still accommodated in the catalytic core of the DNMT1:H3Ub2 complex.

Collectively, these data indicate that the RFTS domain is not completely sequestered from the catalytic domain when H3Ub2 binds to the RFTS domain. However, the structural perturbation between the RFTS and catalytic domains in DNMT1 upon binding of H3Ub2 was enough for the hemimethylated DNA to penetrate into the active center of the catalytic domain.

**Supplementary Note 2. Recognition DNA^mCG/fCG^ by DNMT1 in the ternary complex**

DNA^mCG/fCG^ duplex is embedded into the catalytic core of the DNMT1, and that the 5-fluorocytosine is flipped-out from the DNA duplex (Extended Data Fig. 5b). In addition, the reactive Cys1226 forms a covalent bond with the 6-carbon of 5-fluorocytosine, while the 5-carbon was found to be protruding out from the π-plane of the pyrimidine ring of 5-fluorocytosine (Extended Data Fig. 5b). The cryo-EM map also captures the conjugation step of 5-fluoro substituent by Cys1226, immediately after transfer of the methyl substituent from SAM (Extended Data Fig. 5b). The resultant S-adenosyl-homocysteine (SAH) was observed at 3.4 Å, away from transferred methyl substituent. The catalytic loop containing residues 1224-1238, including the reactive Cys1226, is inserted into the DNA minor groove at the mCG/fCG site. Arg1234 recognizes the 2-carbonyl group of 5-methlcytosine, while the CH-π cluster residues (Arg1234-Phe1235-Arg1238) pushes out the 5-methlcytosine from the major groove (Extended Data Fig. 5c,d). The exposed methyl group of 5-methylcytosine in DNA^mCG/fCG^ duplex is surrounded by the hydrophobic residues, Cys1499, Leu1500, Trp1510, Leu1513 and Met1533 of the TRD (Extended Data Fig. 5c). Recognition of mCG/fCG sequence by the catalytic loop and the TRD is similar to that in the previously reported DNMT1:DNA^mCG/fCG^ structures ^5–7^.

**Supplementary Fig. 1.**
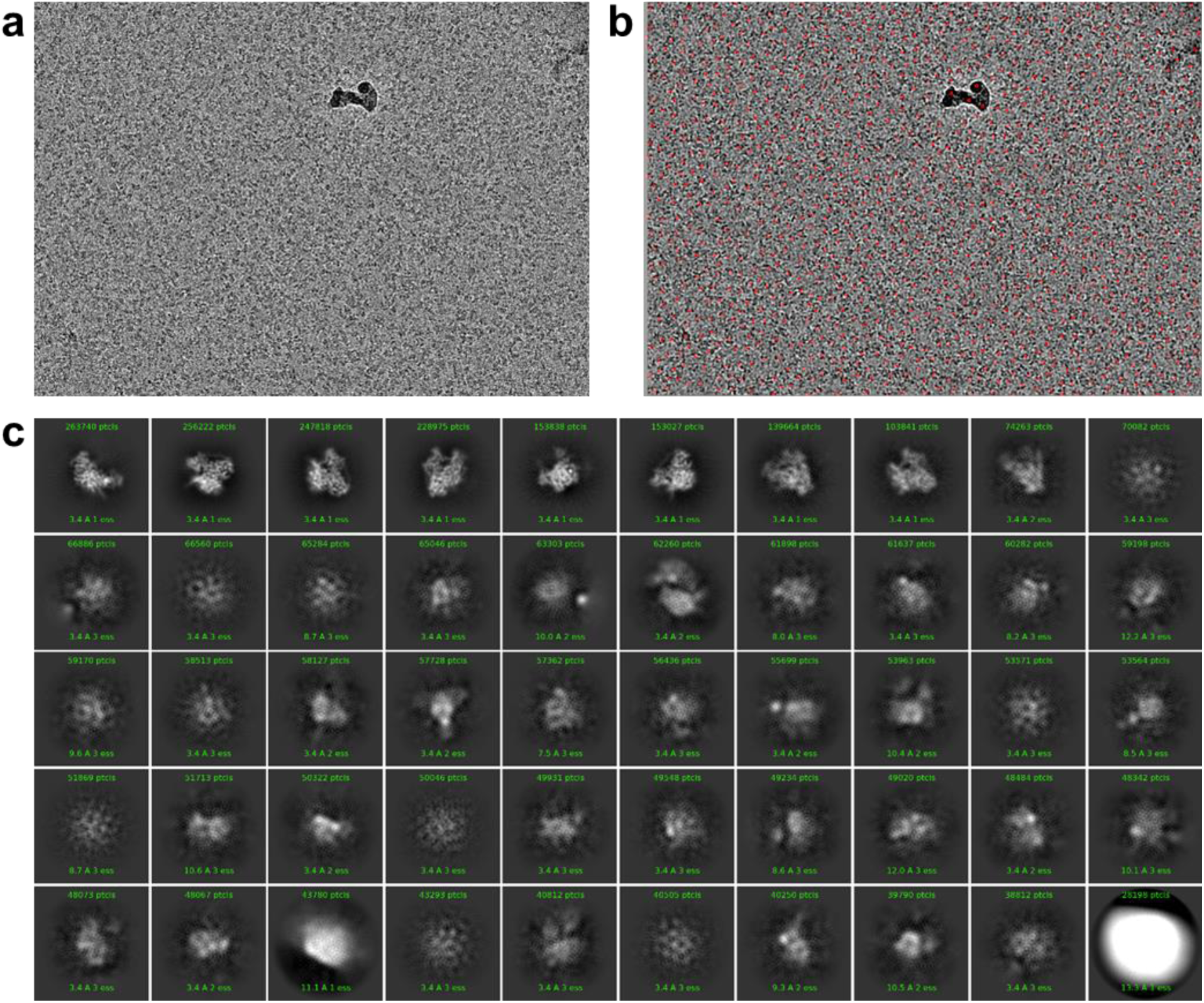
Cryo-EM images of DNMT1:H3Ub2:DNA^mCG/fCG^. **(a)** Motion corrected example of cryo-EM image. **(b)** Initial picked particles are marked with red dot. **(c)** Initial reference-free 2D class averages of the DNMT1:H3Ub2:DNA^mCG/fCG^ ternary complex.

**Supplementary Fig. 2.**
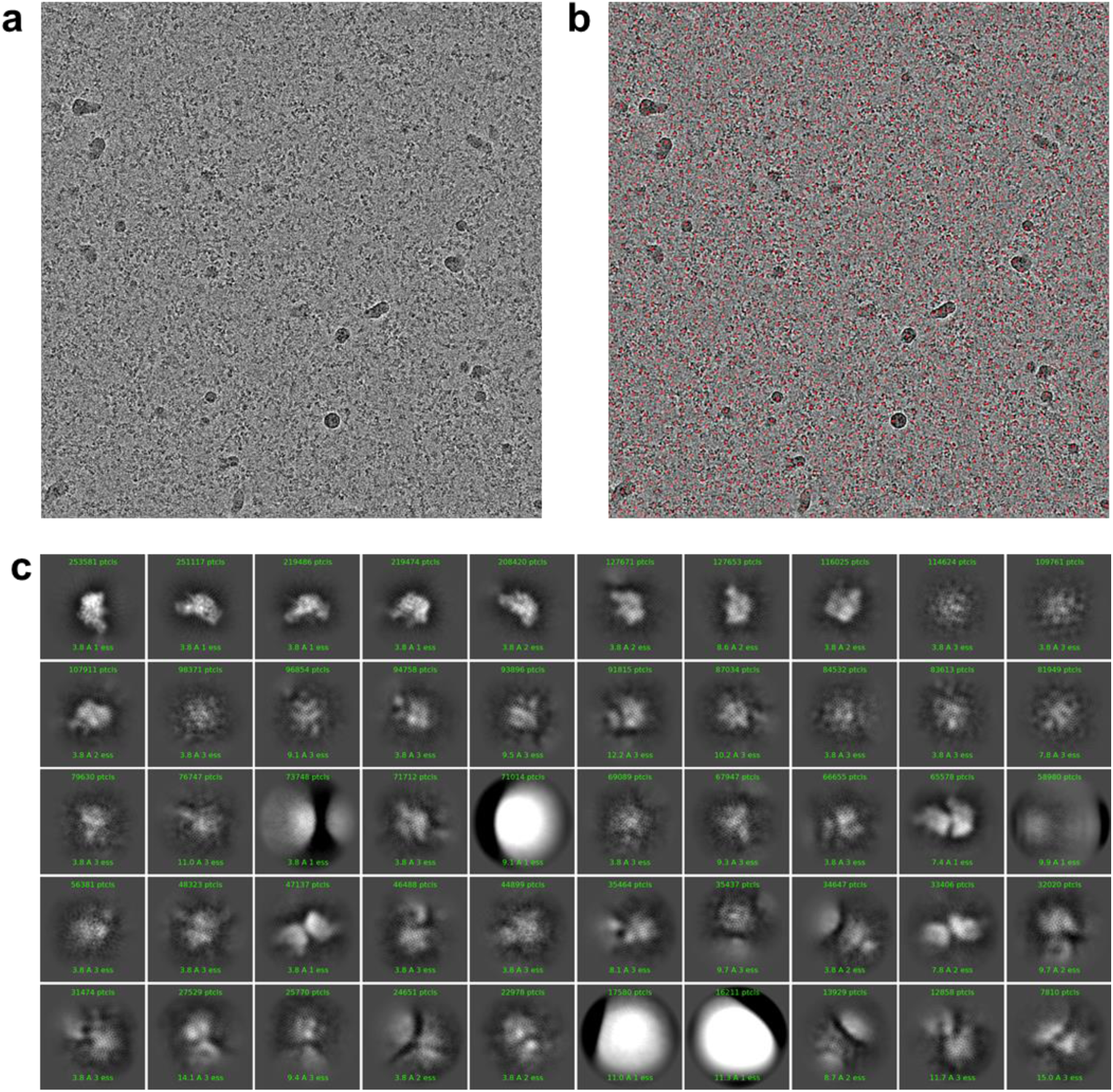
Cryo-EM images of apo-DNMT1. **(a)** Motion corrected example of cryo-EM image. **(b)** Initial picked particles are marked with red dot. **(c)** Initial reference-free 2D class averages of the ternary complex.

**Supplementary Fig. 3.**
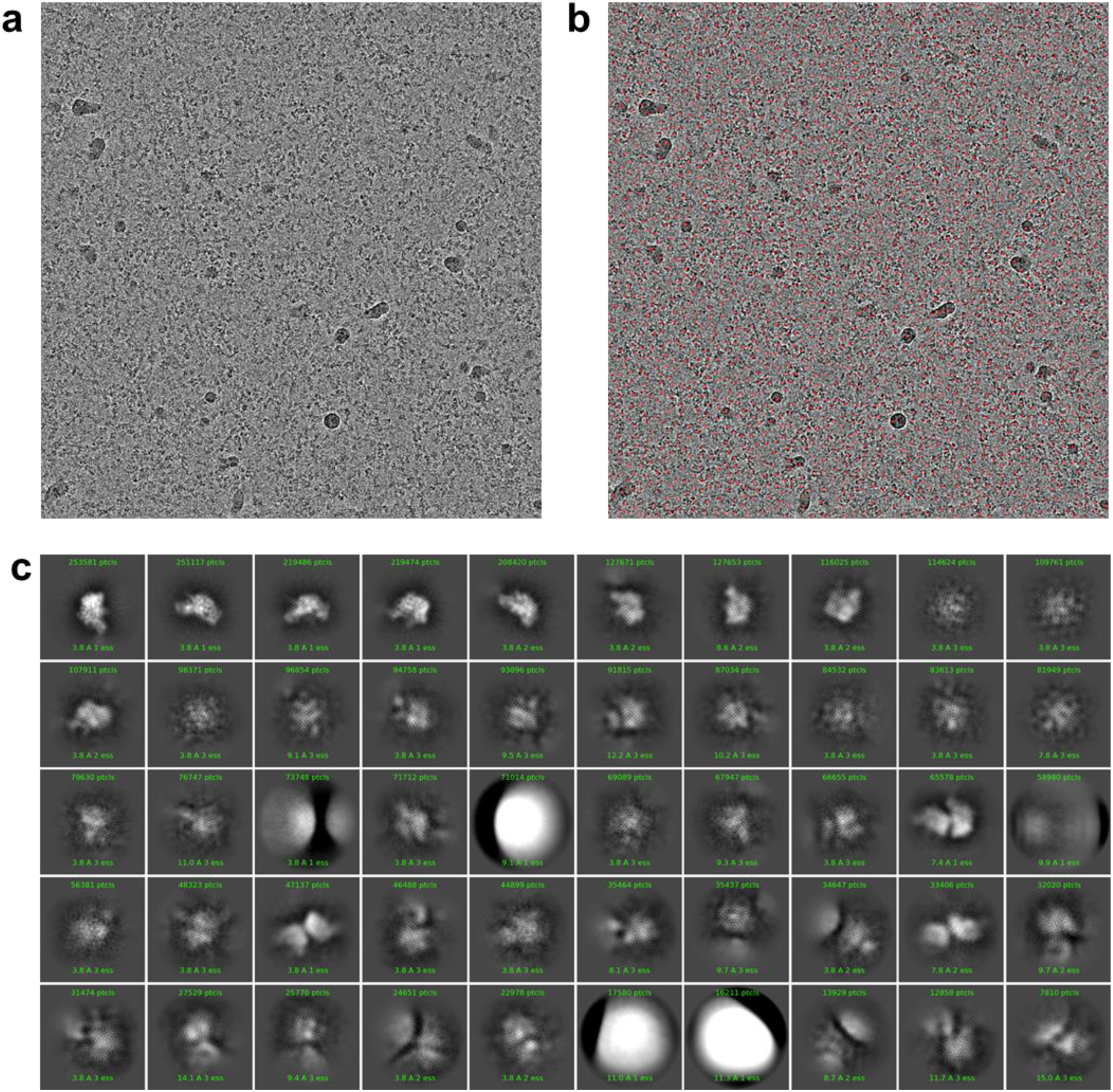
Cryo-EM images of DNMT1:H3Ub2. **(a)** Motion corrected example of cryo-EM image. **(b)** Initial picked particles are marked with red dot. **(c)** Initial reference-free 2D class averages of the ternary complex.

**Supplementary Fig. 4.**
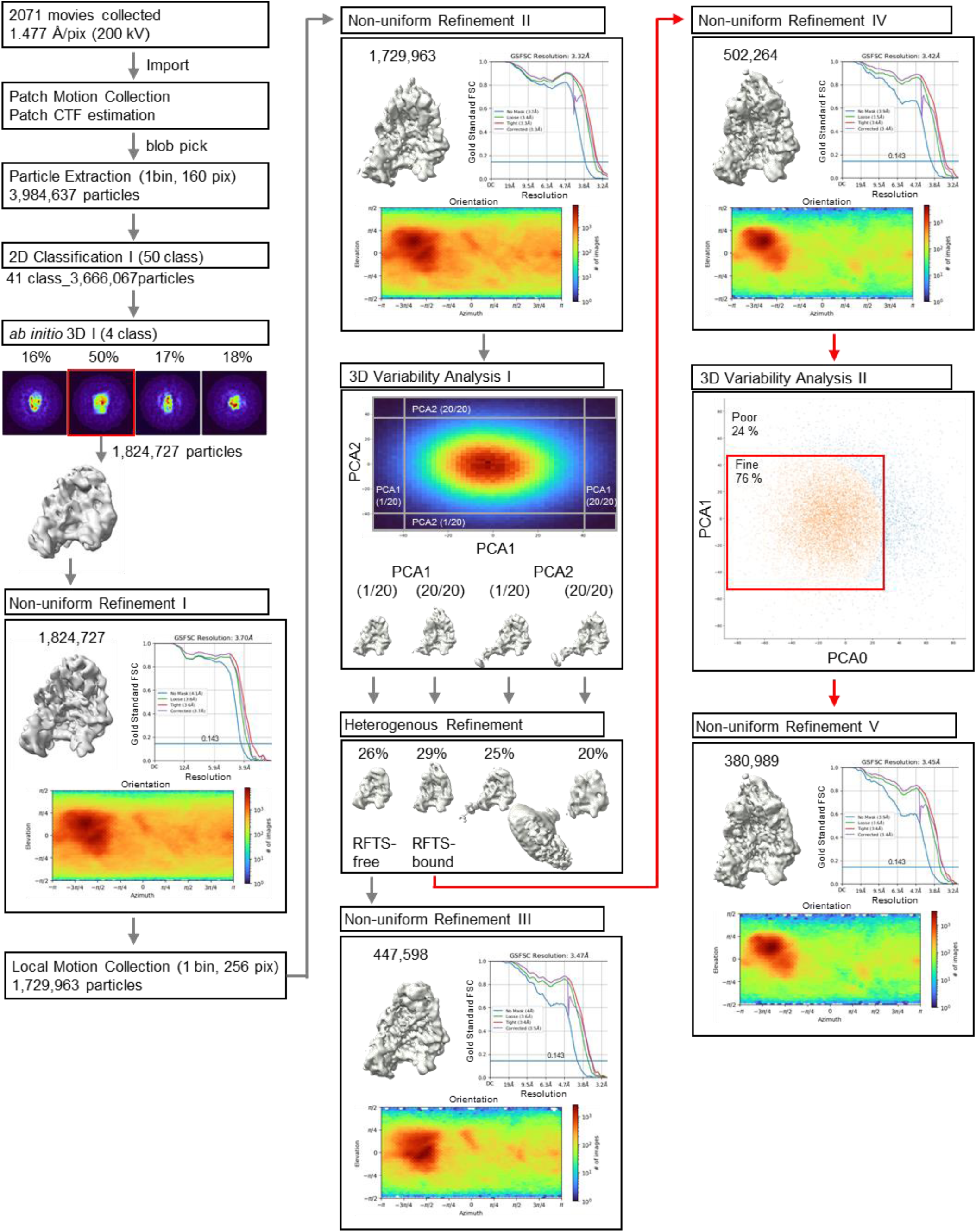
Cryo-EM single particle analysis of apo-DNMT1. Data processing were performed on cryoSPARC. A total of 3,984,637 particles were initially extracted from 2,071 movies with two times binning. After the reference-free 2D classifications, four *ab initio* 3D model were reconstructed from 3,666,067 particles. Non-uniform refinement I was performed against the particles classified in the *ab initio* model of apo-DNMT1 (1,824,727). These particles were re-extracted in a box size of 256 pixels with a 1.477 Å/pixel size by local motion correction. Non-uniform refinement II yields the cryo-EM map with an overall resolution of 3.32 Å resolution. The subsequent initial 3D variability refinement and heterogeneous refinement separated to 477,598 particles as a RFTS-free class and 380,989 particles as a RFTS-bound class with no dimer particle. Finally, 3D variability refinement was performed to remove poor images. The particles (380,989) were then subjected to non-uniform refinement V to yield final cryo-EM map. Overall resolution was estimated to 3.45 Å resolution using the gold-standard Fourier shell correlation with a 0.143 cut-off.

**Supplementary Fig. 5.**
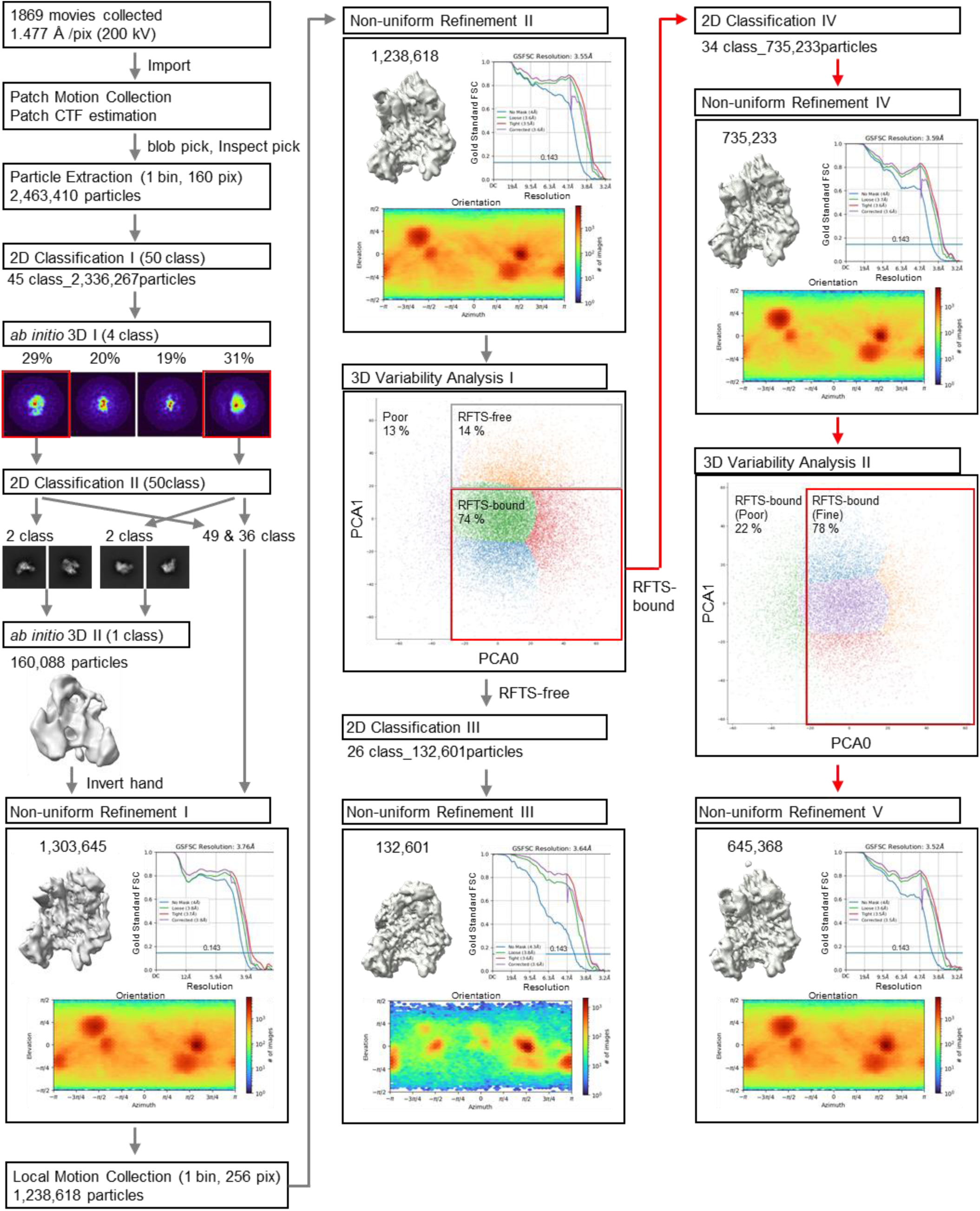
Cryo-EM single particle analysis of DNMT1:H3Ub2. Data processing were performed on cryoSPARC. A total of 2,463,410 particles were initially extracted from 1,869 movies with two times binning. After the reference-free 2D classifications, four *ab initio* 3D model were reconstructed from 2,336,267 particles. The additional 2D classifications were performed for each *ab initio* 3D class, individually. The best four 2D clusters with different orientation were selected for the reconstruction of accurate 3D model. The hand flipped *ab initio* 3D model and all particles without ice template were used for Non-uniform refinement I. These particles were re-extracted in a box size of 256 pixels by local motion correction. Non-uniform refinement II yields the cryo-EM map with an overall resolution of 3.554 Å resolution. The subsequent initial 3D variability refinement separated to 132,601 particles as a RFTS-free class and 753,233 particles as a RFTS-bound class with no dimer particle. Finally, 3D variability refinement was performed to remove poor images. The particles (645,368) were then subjected to non-uniform refinement V to yield final cryo-EM map. Overall resolution was estimated to 3.52 Å resolution using the gold-standard Fourier shell correlation with a 0.143 cut-off.

**Supplementary Fig. 6.**
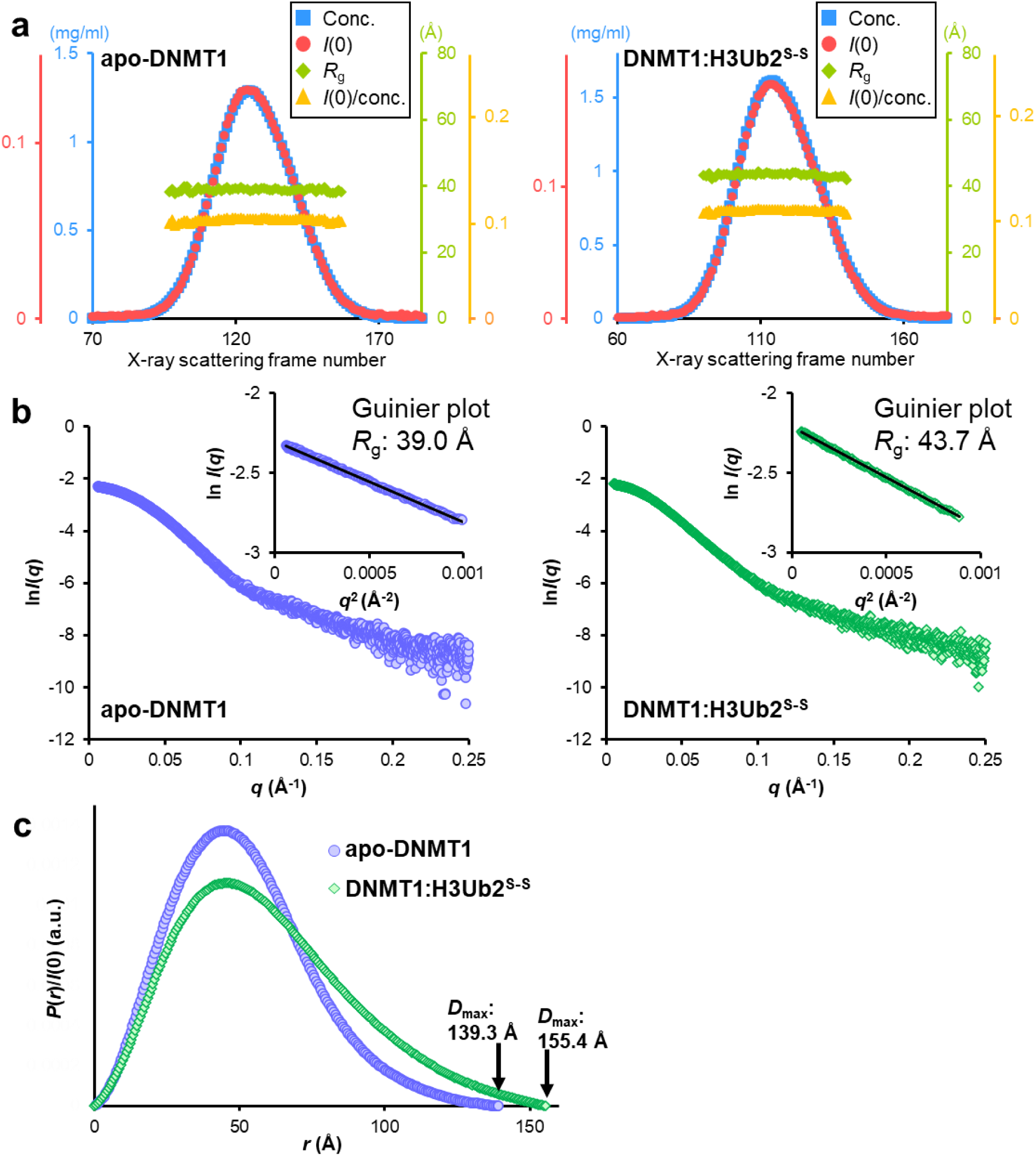
SAXS analysis. **(a)** Sample concentration (blue), *I*(0) (red), *R*_g_ (green) and *I*(0)/*c* (yellow) plots for SEC-SAXS of apo-DNMT1 (left) and DNMT1:H3Ub2^S-S^ (right). **(b)** Experimental X-ray scattering curves of apo-DNMT1 (left) and DNMT1:H3Ub2^S-S^ (right). Vertical and horizontal axes indicate absolute intensity ln*I*(*q*) and scattering angle *q* = 4πsinθ/λ, respectively. Inset indicates Guinier plot with showing *R*_g_ value. **(c)** Pair distance distribution functions *P*(*r*) of apo-DNMT1 (purple) and DNMT1:H3Ub2^S-S^ (green) determined from SAXS data. The *P*(*r*) functions were normalized by *I*(0) calculated from each scatter plot.

## Supplementary Tables

**Supplementary Table 1.** Cryo-EM data collection and refinement statics of apo-DNMT1 and DNMT1:H3Ub2 binary complex.

|  | **apo-DNMT1** | **DNMT1:H3Ub2<sup>iso</sup>** | **Source** |
| --- | --- | --- | --- |
| EMDB number | EMD-33299 | EMD-33298 |  |
| **Data-collection parameters** |  |  |  |
| Microscope | Tecnai Arctica |  |  |
| Detector | K2 |  |  |
| Magnification | 23,500 × |  |  |
| Voltage (kV) | 200 |  |  |
| Electron expose/frame (e<sup>-</sup>/Å<sup>2</sup>) | 1.25 |  |  |
| Defocus range (μm) | 0.8-1.4 |  |  |
| Pixel size (Å) | 1.477 |  |  |
| Number of images | 2,071 | 1,869 |  |
| Initial particle images | 3,984,637 | 2,463,410 | cryoSPARC |
| Final particle images | 380,989 | 645,368 | cryoSPARC |
| Map Resolution (Å) global | 3.45 | 3.52 | cryoSPARC |
| FSC threshold | 0.143 |  |  |
| Symmetry imposed | C1 |  |  |

**Supplementary Table 2.**
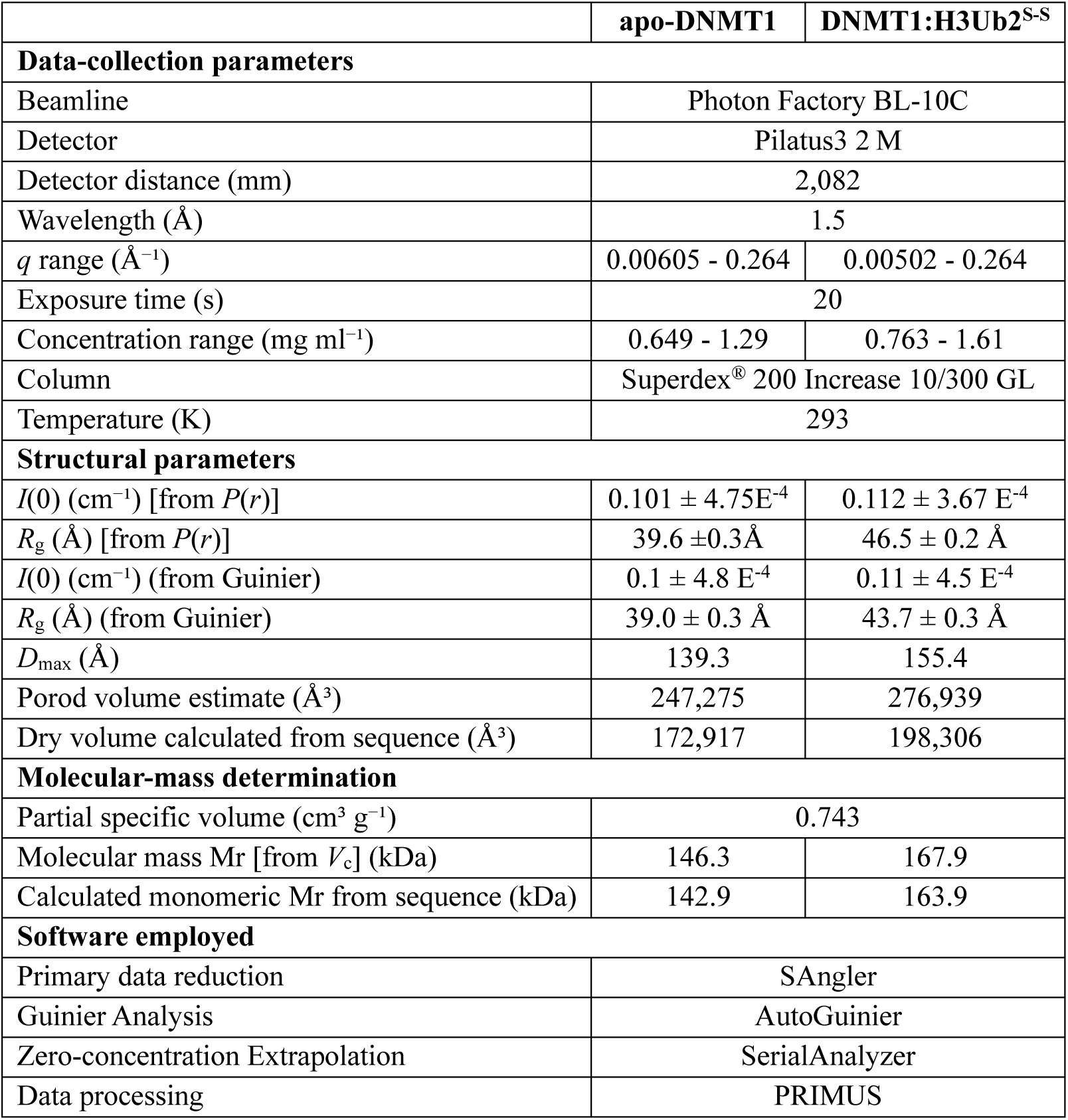
SAXS data collection and scattering derived parameters of apo-DNMT1 and DNMT1:H3Ub2 binary complex.

## Notes

### Competing Interest Statement

The authors have declared no competing interest.

### Summary of Updates

The revised manuscript includes the some changes according to the reviewer's suggestions.

## References

1. Greenberg, M. V. C. & Bourc’his, D. The diverse roles of DNA methylation in mammalian development and disease. Nature reviews. Molecular cell biology 20, 590– 607 (2019).

2. Schübeler, D. Function and information content of DNA methylation. Nature 517, 321– 326 (2015).

3. Nishiyama, A. & Nakanishi, M. Navigating the DNA methylation landscape of cancer. Trends in Genetics 37, 1012–1027 (2021).

4. Petryk, N., Bultmann, S., Bartke, T. & Defossez, P. A. Staying true to yourself: mechanisms of DNA methylation maintenance in mammals. Nucleic acids research 49, 3020–3032 (2021).

5. Bostick, M. et al. UHRF1 plays a role in maintaining DNA methylation in mammalian cells. Science (New York, N.Y.) 317, 1760–1764 (2007).

6. Sharif, J. et al. The SRA protein Np95 mediates epigenetic inheritance by recruiting Dnmt1 to methylated DNA. Nature 450, 908–912 (2007).

7. Arita, K., Ariyoshi, M., Tochio, H., Nakamura, Y. & Shirakawa, M. Recognition of hemi-methylated DNA by the SRA protein UHRF1 by a base-flipping mechanism. Nature 455, 818–821 (2008).

8. Hashimoto, H. et al. The SRA domain of UHRF1 flips 5-methylcytosine out of the DNA helix. Nature 455, 826–829 (2008).

9. Avvakumov, G. V et al. Structural basis for recognition of hemi-methylated DNA by the SRA domain of human UHRF1. Nature 455, 822–825 (2008).

10. Qin, W. et al. DNA methylation requires a DNMT1 ubiquitin interacting motif (UIM) and histone ubiquitination. Cell research 25, 911–29 (2015).

11. Nishiyama, A. et al. Uhrf1-dependent H3K23 ubiquitylation couples maintenance DNA methylation and replication. Nature 502, 249–53 (2013).

12. Ishiyama, S. et al. Structure of the Dnmt1 Reader Module Complexed with a Unique Two-Mono-Ubiquitin Mark on Histone H3 Reveals the Basis for DNA Methylation Maintenance. Molecular Cell 68, 350–360.e7 (2017).

13. Ren, W. et al. Direct readout of heterochromatic H3K9me3 regulates DNMT1-mediated maintenance DNA methylation. Proceedings of the National Academy of Sciences of the United States of America 117, 18439–18447 (2020).

14. Karg, E. et al. Ubiquitome Analysis Reveals PCNA-Associated Factor 15 (PAF15) as a Specific Ubiquitination Target of UHRF1 in Embryonic Stem Cells. Journal of molecular biology 429, 3814–3824 (2017).

15. Ren, W., Gao, L. & Song, J. Structural Basis of DNMT1 and DNMT3A-Mediated DNA Methylation. Genes 9, (2018).

16. Zhang, Z. M. et al. Crystal Structure of Human DNA Methyltransferase 1. Journal of molecular biology 427, 2520–2531 (2015).

17. Takeshita, K. et al. Structural insight into maintenance methylation by mouse DNA methyltransferase 1 (Dnmt1). Proceedings of the National Academy of Sciences of the United States of America 108, 9055–9059 (2011).

18. Song, J., Teplova, M., Ishibe-Murakami, S. & Patel, D. J. Structure-based mechanistic insights into DNMT1-mediated maintenance DNA methylation. Science 335, 709–712 (2012).

19. Adam, S. et al. DNA sequence-dependent activity and base flipping mechanisms of DNMT1 regulate genome-wide DNA methylation. Nature Communications 2020 11:1 11, 1–15 (2020).

20. Pappalardi, M. B. et al. Discovery of a first-in-class reversible DNMT1-selective inhibitor with improved tolerability and efficacy in acute myeloid leukemia. Nature Cancer 2021 2:10 2, 1002–1017 (2021).

21. Song, J., Rechkoblit, O., Bestor, T. H. & Patel, D. J. Structure of DNMT1-DNA complex reveals a role for autoinhibition in maintenance DNA methylation. Science 331, 1036– 1040 (2011).

22. Osterman, D. G., Depillis, G. D., Wu, J. C., Matsuda, A. & Santi, D. V. 5-Fluorocytosine in DNA is a mechanism-based inhibitor of HhaI methylase. Biochemistry 27, 5204–5210 (1988).

23. Punjani, A. & Fleet, D. J. 3D variability analysis: Resolving continuous flexibility and discrete heterogeneity from single particle cryo-EM. Journal of structural biology 213, (2021).

24. Nishiyama, A. et al. Two distinct modes of DNMT1 recruitment ensure stable maintenance DNA methylation. Nature Communications 11, 1222 (2020).

25. Jia, D., Jurkowska, R. Z., Zhang, X., Jeltsch, A. & Cheng, X. Structure of Dnmt3a bound to Dnmt3L suggests a model for de novo DNA methylation. Nature 449, 248–251 (2007).

26. Lin, C. C., Chen, Y. P., Yang, W. Z., Shen, J. C. K. & Yuan, H. S. Structural insights into CpG-specific DNA methylation by human DNA methyltransferase 3B. Nucleic acids research 48, 3949–3961 (2020).

27. Zhang, Z. M. et al. Structural basis for DNMT3A-mediated de novo DNA methylation. Nature 554, 387–391 (2018).

28. Gao, L. et al. Comprehensive structure-function characterization of DNMT3B and DNMT3A reveals distinctive de novo DNA methylation mechanisms. Nature Communications 2020 11:1 11, 1–14 (2020).

29. Punjani, A., Rubinstein, J. L., Fleet, D. J. & Brubaker, M. A. cryoSPARC: algorithms for rapid unsupervised cryo-EM structure determination. Nature methods 14, 290–296 (2017).

30. Frauer, C. et al. Recognition of 5-hydroxymethylcytosine by the Uhrf1 SRA domain Supplementary Table S1. Sequences of DNA oligonucleotides used for preparation of double stranded fluorescent DNA substrates.

31. Scheres, S. H. W. & IUCr. Amyloid structure determination in RELION-3.1. urn:issn:2059-7983 76, 94–101 (2020).

32. Pettersen, E. F. et al. UCSF ChimeraX: Structure visualization for researchers, educators, and developers. Protein science : a publication of the Protein Society 30, 70–82 (2021).

33. Shimizu, N., et al. Software development for analysis of small-angle x-ray scattering data. in AIP Conference Proceedings 1741, 050017 (AIP Publishing LLC, 2016).

34. Orthaber, D., Bergmann, A. & Glatter, O. SAXS experiments on absolute scale with Kratky systems using water as a secondary standard. Journal of Applied Crystallography 33, 218–225 (2000).

35. Yonezawa, K., Takahashi, M., Yatabe, K., Nagatani, Y. & Shimizu, N. Software for serial data analysis measured by SEC-SAXS/UV-Vis spectroscopy. in 060082 (2019). doi:10.1063/1.5084713

36. Rambo, R. P. & Tainer, J. A. Accurate assessment of mass, models and resolution by small-angle scattering. Nature 496, 477–481 (2013).

37. Svergun, D. I. Restoring low resolution structure of biological macromolecules from solution scattering using simulated annealing. Biophysical journal 76, 2879–86 (1999).

38. Svergun, D., Barberato, C. & Koch, M. H. CRYSOL – a Program to Evaluate X-ray Solution Scattering of Biological Macromolecules from Atomic Coordinates. urn:issn:0021-8898 28, 768–773 (1995).

39. Kumamoto, S. et al. HPF1-dependent PARP activation promotes LIG3-XRCC1-mediated backup pathway of Okazaki fragment ligation. Nucleic acids research 49, 5003–5016 (2021).

40. Yesbolatova, A., Natsume, T., Hayashi, K. ichiro & Kanemaki, M. T. Generation of conditional auxin-inducible degron (AID) cells and tight control of degron-fused proteins using the degradation inhibitor auxinole. Methods (San Diego, Calif.) 164–165, 73–80 (2019).

41. Natsume, T., Kiyomitsu, T., Saga, Y. & Kanemaki, M. T. Rapid Protein Depletion in Human Cells by Auxin-Inducible Degron Tagging with Short Homology Donors. Cell reports 15, 210–218 (2016).

42. Mátés, L. et al. Molecular evolution of a novel hyperactive Sleeping Beauty transposase enables robust stable gene transfer in vertebrates. Nature genetics 41, 753–761 (2009).

## References

1. Ishiyama, S. et al. Structure of the Dnmt1 Reader Module Complexed with a Unique Two-Mono-Ubiquitin Mark on Histone H3 Reveals the Basis for DNA Methylation Maintenance. Molecular Cell 68, 350–360.e7 (2017).

2. Zhang, Z. M. et al. Crystal Structure of Human DNA Methyltransferase 1. Journal of molecular biology 427, 2520–2531 (2015).

3. Takeshita, K. et al. Structural insight into maintenance methylation by mouse DNA methyltransferase 1 (Dnmt1). Proceedings of the National Academy of Sciences of the United States of America 108, 9055–9059 (2011).

4. Syeda, F. et al. The replication focus targeting sequence (RFTS) domain is a DNA-competitive inhibitor of Dnmt1. The Journal of biological chemistry 286, 15344–15351 (2011).

5. Pappalardi, M. B. et al. Discovery of a first-in-class reversible DNMT1-selective inhibitor with improved tolerability and efficacy in acute myeloid leukemia. Nature Cancer 2021 2:10 2, 1002–1017 (2021).

6. Adam, S. et al. DNA sequence-dependent activity and base flipping mechanisms of DNMT1 regulate genome-wide DNA methylation. Nature Communications 2020 11:1 11, 1–15 (2020).

7. Song, J., Teplova, M., Ishibe-Murakami, S. & Patel, D. J. Structure-based mechanistic insights into DNMT1-mediated maintenance DNA methylation. Science 335, 709–712 (2012).

